# Cardiac contraction and relaxation are regulated by beta 1 adrenergic receptor-generated cAMP pools at distinct membrane locations

**DOI:** 10.1101/2022.07.13.499965

**Authors:** Ting-Yu Lin, Quynh N. Mai, Hao Zhang, Emily Wilson, Huan-Chieh Chien, Sook Wah Yee, Kathleen M. Giacomini, Jeffrey E. Olgin, Roshanak Irannejad

**Affiliations:** Cardiovascular Research Institute, University of California, San Francisco, USA; Department of Medicine, Division of Cardiology, University of California San Francisco, San Francisco, USA; Department of Bioengineering and Therapeutic Sciences, University of California San Francisco, California, USA; Institute for Human Genetics, University of California San Francisco, California; Department of Biochemistry & Biophysics, University of California, San Francisco, USA

## Abstract

Cells interpret a variety of signals through G protein-coupled receptors (GPCRs) and stimulate the generation of second messengers such as cyclic adenosine monophosphate (cAMP). A long-standing puzzle is deciphering how GPCRs elicit different physiological responses despite generating similar levels of cAMP. We previously showed that some GPCRs generate cAMP from both the plasma membrane and the Golgi apparatus. Here, we demonstrate that cardiomyocytes distinguish between subcellular cAMP inputs to elicit different physiological outputs. We show that generating cAMP from the Golgi leads to regulation of a specific PKA target that increases the rate of cardiomyocyte relaxation. In contrast, cAMP generation from the plasma membrane activates a different PKA target that increases contractile force. We further validated the physiological consequences of these observations in intact zebrafish and mice. Thus, we demonstrate that the same GPCR acting through the same second messenger regulates cardiac contraction and relaxation dependent on its subcellular location.

## Introduction

GPCRs are the largest family of membrane receptors that communicate downstream signaling pathways to regulate cellular functions. Many of them signal through the stimulatory G protein (Gs) to generate cAMP^1^. There are number of different hormones that stimulate cAMP generation through the activation of different types of GPCRs that are expressed in the same cell^2,3^. However, activation of each GPCR can trigger different physiological responses despite generating the same level of cAMP. A classic example of such distinct physiological responses is the activation of beta-adrenergic receptors (βARs) and prostaglandin E1-type receptors, which trigger similar elevations of cAMP in cardiac tissue, but only the activated βAR causes increased cardiac contractility and glycogen metabolism in cardiomyocytes^4–7^. Analogous observations have been reported even within the same family of receptors. For example, β1AR and β2AR, the two main beta-adrenergic (βAR) family isoforms in cardiomyocytes are both activated by sympathetic hormones epinephrine and norepinephrine, and trigger Gs-mediated cAMP generation^8,9^. Nevertheless, these receptors elicit distinct effects on cardiac function. In healthy cardiomyocytes, β1AR signaling regulates chronotropy (heart rate), inotropy (force of contraction), and lusitropy (relaxation)^10–12^ while β2AR signaling only modestly contributes to chronotropy and has no appreciable effect on lusitropy in murines^13^. In the context of heart failure, β1AR signaling promotes cardiomyocyte hypertrophy and apoptosis, whereas β2AR signaling inhibits both^10,12–16^. Numerous hypotheses have been advanced to explain how β1AR and β2AR function differently from each other, both in health and disease states, but answers have been elusive. Importantly, β1AR and β2AR localize to different subdomains in cardiomyocytes. While β1AR is mostly at the plasma and Golgi membranes, β2AR is mostly localized in transverse-tubules (t-tubule). Whether the distinct βAR localizations underlie the noted physiological and pathophysiological outcomes attributed to each has not been addressed. In the past decade, several reports have shown that GPCR can signal from subcellular membrane compartments^17–25^. For example, we have shown that β1AR can be activated and generate cAMP from the plasma membrane and the Golgi apparatus^19^, whereas β2AR can generate cAMP responses from the plasma membrane and endosomes^26^. The significance of generating cAMP from distinct membrane compartments is beginning to be understood. There is evidence for distinct transcriptional responses^27,28^ and activating distinct signaling pathways depending on the subcellular source of cAMP production^9,27,29–32^.

Classically, cAMP was considered a highly diffusible molecule, and thus it was reasoned that cAMP generation at the plasma membrane, by GPCR/Gs activation, is sufficient to activate downstream effectors of cAMP in other subcellular membrane compartments. Recent reports, however, demonstrate that cAMP is mostly immobile and constrained due to binding to specific cAMP binding sites. Several studies have reported the role of phosphodiesterases (PDEs) and the regulatory subunit of protein kinase A (PKA), the main cAMP binding protein, in constraining cAMP at specific membrane compartments ^33–36^. In the basal state, cAMP was shown to be mostly bound to intracellular cAMP binding sites, such as PKA regulatory subunits, at each subcellular location and PDEs can generate a nanometer-size domain around a source of cAMP^37,38^. Moreover, the PKA regulatory subunit has been shown to form a liquid-liquid phase in the cytoplasm and sequester cAMP, thereby acting as a sponge to buffer cAMP in the cytoplasm^39^. It is only after the elevation of cAMP in cells, that free cAMP can act on PKA and other effectors to initiate downstream cellular responses. Furthermore, the catalytic activity of PKA has also been shown to be constrained to targets within a radius of 15-25 nm^40^. How this spatially and functionally restricted PKA can phosphorylate downstream targets localized within the cells is unclear. Thus, the nanometer scale of the cAMP diffusion range conflicts with the prevalent model whereby plasma membrane-localized receptors generate cAMP which then propagates linearly to activate cAMP-mediated PKA responses in distant subcellular locations^36,41,42^.

The present study reveals the pivotal role of local generation of cAMP in controlling local PKA activation at specific subcellular compartments. We demonstrate how cells with more complex architecture such as cardiomyocytes can precisely sense subcellular cAMP pools and regulate local PKA activity to generate compartment-specific cellular and physiological outputs.

To determine the relevance of local cAMP generation and the activity map of cAMP around activated β1AR at distinct membrane locations, we measured the activation of downstream effectors of cAMP/PKA in cardiomyocytes. Using an optogenetic approach, we show that local generation of cAMP at the Golgi leads to distinct activation of downstream effector of PKA that increases the rate of relaxation in cardiomyocytes. Conversely, we demonstrated that activation of the plasma membrane pool of β1AR, using pharmacological and genetic approaches, leads to the activation of proximal PKA effectoβrs at the plasma membrane that increase the force of contraction in cardiomyocytes. Finally, we tested two different animal models, zebrafish and mice, using optogenetic and pharmacological approaches and found distinct regulation of cardiac inotropy and lusitropy by different cAMP pools.

## Result

### An optogenetic system to generate cAMP at the Golgi membrane in cardiomyocytes

To assess whether cAMP generation from the Golgi membrane communicates different cellular information, we developed an optogenetic system based on a bacterial photo-activatable adenylyl cyclase (bPAC). bPAC had been previously used to generate cAMP from distinct cellular compartments such as endosomes and cilia^28,43^. We fused bPAC to the trans Golgi Network 46 protein, a known Golgi-targeting motif, to target bPAC to the trans Golgi membrane (Extended Data Fig. 1a). We also confirmed Golgi localization in cardiomyocytes (Fig. 1b and c). To assess whether cardiomyocytes expressing Golgi-bPAC generate cAMP in response to blue light treatment, we virally transduced neonatal cardiomyocytes and measured cAMP concentrations. Treating the cells with 0.34 μW/cm^2^ blue light for 3 minutes resulted in ~20 pmol/mg cAMP accumulation in cardiomyocytes (Fig. 1c). This is consistent with the physiological levels of cAMP in cardiomyocytes in response to β1AR stimulation with 100 nM epinephrine (Extended Data Fig. 1c). Considering that the average volume of cardiomyocytes is reported at around 15 picolitres (15000 μm^3^)^44^, this concentration translates into ~ 1 μM cAMP in each cell, which is within the physiological levels of cAMP upon hormone stimulation^37,45^.

**Figure 1.**
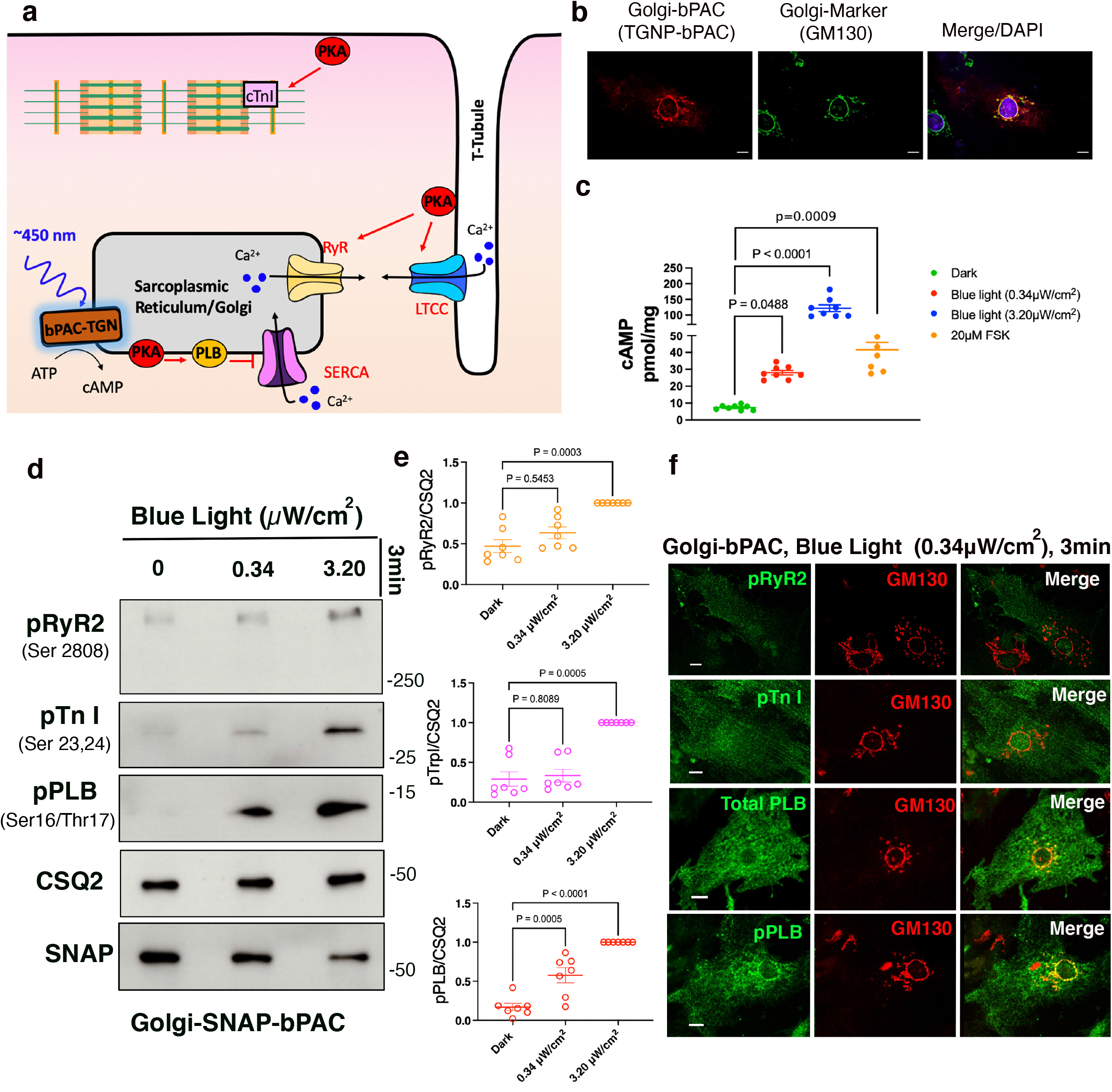
cAMP generation at the Golgi distinctly regulate cardiomyocyte relaxation in neonatal mouse cardiomyocytes. **a**. Illustration of the roles of subcellular cAMP/PKA signaling hubs in regulating contraction/relaxation responses of cardiomyocytes. The cAMP/PKA signaling hub on the sarcolemma and t-tubule increase cardiac muscle contraction by promoting the phosphorylation of cardiac troponin I (cTnI), ryanodine receptor 2 (RyR2), and L-type calcium channel (LTCC). Activation of the Golgi pool of cAMP/PKA increases the phospholamban (PLB) phosphorylation to induce cardiac muscle relaxation by promoting sarcoplasmic reticulum Ca^2+^-ATPase (SERCA)-mediated calcium uptake. After blue light (~450 nm) stimulation, Golgi-targeting bPAC (TGNP-bPAC) produces cAMP to activate cAMP/PKA signaling near the Golgi membranes. **b**. Golgi-bPAC was generated by fusing the bPAC protein to the transmembrane and cytoplasmic domain of TGNP with a SNAP-tag. Representative images of Golgi-bPAC (red) and Golgi marker (green), visualized by SNAP and GM130 antibodies, and DAPI staining. Scale bar, 10 *μ*m. **c**. cAMP generation mediated by Golgi-bPAC in neonatal cardiomyocytes. Cells were stimulated with blue light for 3 minutes and 5 minutes or forskolin (FSK 20μM) for 5 min and then lysed for direct cAMP determination by ELISA. cAMP concentrations were normalized to the relative protein concentrations in cell lysate of each sample. The quantified data are represented as mean ± S.E.M. The p-values were calculated by one-way ANOVA. n=8 and 6 biological replicates form Golgi-bPAC and FSK treated cardiomyocytes, respectively. **d**. Representative phosphorylation profiles of RyR2, TnI, and PLB induced by Golgi-bPAC in mouse neonatal cardiomyocytes. The protein levels of p-RyR2 Ser2808, p-TnI Ser23/24, and p-PLB Ser16/Thr17 were analyzed in the Golgi-bPAC-expressing mouse neonatal cardiomyocytes kept in the dark or exposed to 0.34 or 3.2 *μ*W/cm^2^ blue light for 3 minutes. The protein level of Golgi-bPAC was analyzed using the SNAP antibody. The protein level of CSQ2 was used as a loading control. **e**. The band intensities of p-RyR2, p-TnI, and p-PLB were normalized to CSQ2 intensity and then reported as a percentage of the highest value in the groups. The quantified data from different experiments are presented as mean ± S.E.M. The p-values were calculated by one-way ANOVA. n=7 biological replicates. **f**. The subcellular localization of p-RyR2, p-TnI, PLB, and p-PLB upon stimulation of Golgi-bPAC with 0.34 *μ*W/cm^2^ blue light in mouse neonatal cardiomyocytes. The protein localizations of p-RyR2 Ser2808, p-TnI Ser23/24, total PLB, and pPLB Ser16/Thr17 antibody were visualized (green) with Golgi marker stained by GM130 (red). Scale bar, 10 *μ*m.

Since cAMP generated from activated β1AR results in the activation of PKA in cardiomyocytes, we investigated whether cAMP generation from the Golgi membrane can activate downstream targets of PKA. In cardiomyocytes, β1AR-mediated cAMP generation regulates chronotropy (heart rate), inotropy (force of contraction), and lusitropy (relaxation) through PKA-mediated phosphorylation of proteins, such as cardiac Troponin I (TnI), Ryanodine 2 receptors (RyR2), and phospholamban (PLB) (Fig. 1a)^10^. Thus, we examined whether generating physiological levels of cAMP by Golgi-bPAC can phosphorylate and activate downstream targets of PKA. Golgi-bPAC expression in the absence of blue light had minimal effect on the phosphorylation of PKA effectors (Fig. 1d, left lanes). Notably, 0.34 μW/cm^2^ blue light treatment resulted in the robust phosphorylation of PLB but not TnI and RyR2 (Fig. 1d middle lane and quantification in e). This suggests that cardiomyocytes selectively respond to physiologically relevant Golgi-generated cAMP levels. Consistent with this, subcellularly localized PDEs have been shown to constrain cAMP levels to the vicinity of its site of generation^33–37^.

We then explored whether supraphysiological cAMP levels can overcome this selective response to Golgi-generated cAMP. We manipulated two variables, blue light intensity and exposure time, to reach cAMP levels that are similar to physiological and supraphysiological levels. By increasing blue light intensity to 3.20 μW/cm^2^ for 3 minutes, we found that Golgi-generated cAMP reaches supraphysiological levels and causes the phosphorylation of all three PKA effectors (Fig. 1c and 1d, right lanes and quantification in e). We then tested the consequences of keeping the blue light intensity low (0.34 μW/cm^2^) but increasing the exposure time. As in the case of increased intensity, a 7-minute exposure time also led to the phosphorylation of all effectors (Extended Data Fig. 1b). We further confirmed that phosphorylation of PLB, TnI and RyR2 by Golgi-bPAC is mediated by PKA, as inhibiting PKA activity using PKA inhibitor (H89), diminished phosphorylation of all three effectors (Extended Data Fig. 1e). These results suggest that differential consequences of cAMP generated at the Golgi is mediated by PKA, and that at physiological levels, only PLB, a regulator of lusitropy, is phosphorylated.

PLB is a sarco/endoplasmic reticulum (SR)-localized protein and is the dominant regulator of Ca^2+^ reuptake by sarco/endoplasmic reticulum Ca^2+^ ATPases (SERCA)^46^. We therefore asked whether the local pool of cAMP that is generated by Golgi-bPAC phosphorylates PLB in the vicinity of the Golgi membrane. Immunofluorescence imaging of the non-phosphorylated form of PLB in neonatal cardiomyocytes revealed a SR-localization pattern throughout the cytoplasm, as expected (Fig. 1f, third row). The fluorescent signals were not detectable when we used antibody against phosphorylated form of PLB in cardiomyocytes that were unstimulated by blue light (Figure S1D, top row). When cardiomyocytes were stimulated with 0.34 μW/cm^2^ blue light for 3 minutes, immunofluorescence imaging of phosphorylated PLB (pPLB) detected distribution of pPLB in the vicinity of the Golgi membranes (Fig. 1f, bottom row). The fluorescent signals were not detectable when we used antibodies against the phosphorylated forms of Tn-I and RyR2 at this blue light regimen (Fig. 1f, top rows). However, as we increased cAMP generation by 3.20 μW/cm^2^ blue light stimulation for 3 minutes, phosphorylated forms of all three effectors were detected (Extended Data Fig. 1d). Importantly, at this higher cAMP concentration, pPLB no longer had a near Golgi distribution but showed a broad SR-localization pattern (Extended Data Fig. 1d, second row). Together, these data suggest that cAMP generation from the Golgi membranes at similar levels as those generated by sympathetic hormones results in PKA-mediated phosphorylation of downstream target, PLB, in the vicinity of the Golgi membranes.

### Golgi-cAMP specifically regulates cardiac relaxation in zebrafish

As PLB is the key regulator of cardiomyocyte relaxation (lusitropy), we predicted that cAMP generation at the Golgi specifically regulates lusitropy. To test this hypothesis, we generated a Golgi-bPAC expressing transgenic zebrafish. Zebrafish is a well-established animal model for exploring the physiological parameters of cardiac function. The molecular mechanisms underlying their heart function are very similar to those of higher vertebrates^47^. The optical clarity of zebrafish embryos allows the real-time and *in vivo* visualization of the heart contractility responses and makes it a useful vertebrate model system for studying cardiovascular performance using optogenetic tools ^48,49^. To measure cardiac outputs such as chronotropy, inotropy, and lusitropy, we generated transgenic zebrafish that express Golgi-bPAC (Extended Data Fig. 2a). These transgenic zebrafish were developed from the established line, Tg(Flk-Ras-cherry)^s896^, which expresses Ras-Cherry in the inner layer of the heart wall (endocardium)^50^. As a result, we were able to trace the motion of the walls of the cardiac atrium (A) and ventricle (V) in different phases of the cardiac cycle in the RFP channel using a confocal microscope imaging mCherry (Fig. 2a and b). The coupling of ventricular and atrial contraction can be determined by evaluating the time delay between the peak values of the extracted synchronous chronologies within the same cardiac cycle (Fig. 2b). We were also able to measure cardiac rhythm (heart rate) by measuring the distance between two consecutive highest points of the peaks of each cycle.

**Figure 2.**
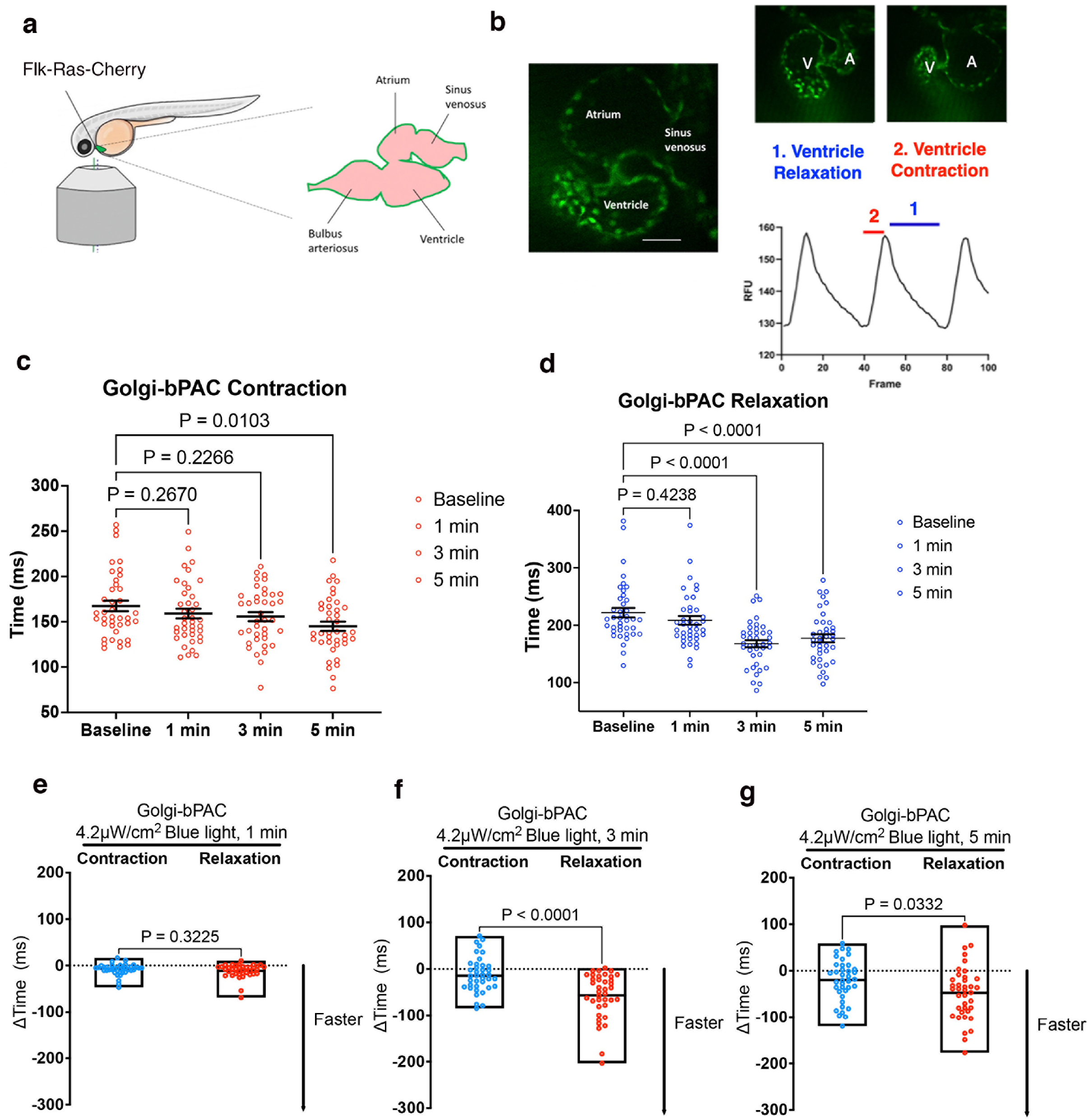
Golgi-delimited cAMP generation promotes faster ventricular relaxation in zebrafish. **a**. Illustration of the mounting position of the zebrafish to image the heart and diagram of the zebrafish heart. **b**. Representative image of a live zebrafish heart (left) and demonstrating ventricular contraction and relaxation in zebrafish heart. Fluctuations in the fluorescence of the heart during contraction and relaxation over time are measured. The time between fluorescence maxima to minima is the time of contraction, and the subsequent fluorescence minima to maxima portion of the graph is measured as the time of relaxation. Sale bar, 50 μm. A, Atrium. V, Ventricle. **c**. Changes of heart contraction and **d**. relaxation time relative to baseline in Golgi-bPAC expressing zebrafish. Basal images of the zebrafish hearts were acquired for 1500 frames. The transgenic Golgi-bPAC zebrafish were exposed to 4.2 *μ*W/cm^2^ blue light and imaged after 1, 3, and 5 minutes stimulation. **e-g**. Comparisons of rates of contraction and relaxation at 1, 3 and 5 minutes after 4.2 *μ*W/cm^2^ blue light illumination. The differences between faster contraction versus relaxation time over baseline are quantified and presented here as mean ± S.E.M. with p-values presented. The floating bars indicate minimum and maximum with line at mean. Data were analyzed by t-test. n=38 zebrafish.

Treating Golgi-bPAC expressing zebrafish with 4.2 μW/cm^2^ blue light for 3 minutes resulted in detectable levels of cAMP in zebrafish (Extended Data Fig. 2b). To test the effect of cAMP accumulation in zebrafish hearts, we first measured their heart rate in 72 hours post fertilization (hpf). A basal heart rate of zebrafish is around 120-180 beats per minute (bpm). Illuminating Golgi-bPAC zebrafish with 4.2 μW/cm^2^ blue light resulted in an increase in the heart rate in a time-dependent manner (Extended Data Fig. 2c). Because the heart rate measurement using this assay is determined by adding the rate of contraction to the rate of relaxation, we sought to specifically evaluate each rate at a given time upon blue light illumination. Treating Golgi-bPAC expressing zebrafish with 4.2 μW/cm^2^ blue light for 1 minutes did not generate measurable cAMP level and had no significant effect on the heart rate, the mean rate of relaxation (lusitropy) or the contraction (inotropy) (Fig. 2c-e). Increasing the blue light illumination time to 3 minutes also did not result in significant change in the mean rate of contraction but increased the mean rate of relaxation (Fig. 2 c and d). Illuminating blue light for 5 minutes resulted in a significant change in the mean rate of contraction (inotropy) (Fig. 2c) and relaxation (lusitropy) (Fig 2d). Importantly, the rate of relaxation increased more significantly when compared to the rate of contraction time at 3 minutes (Fig. 2f). This result is consistent with the specific phosphorylation of PLB but not TnI and RyR2 in cardiomyocytes using low-level blue light treatment. However, at 5 minutes blue light illumination, the rate of relaxation and contraction increased similarly (Fig. 2g).

Treating Golgi-bPAC zebrafish with 4.2 μW/cm^2^ blue light and the PDE inhibitor, 3-isobutyl-1-methylxanthine (IBMX), to increase cAMP levels, elevated the rate of both contraction and relaxation at all time points (Extended Data Fig. 2e). Importantly, stimulating control zebrafish (Tg(Flk-Ras-cherry)^s896^) with 4.2 μW/cm^2^ blue light at 1,3 and 5 minutes did not result in significant changes in the rate of contraction or relaxation (Extended Data Fig. 2d). Altogether, these results suggest that cAMP generation from the Golgi at physiological levels specifically regulates the rate of cardiac relaxation responses. However, this differential interpretation of cAMP is disrupted when cAMP generation at the Golgi is increased to supraphysiological levels and PDEs are no longer able to constrain cAMP to the vicinity of a compartment^33–36^.

### Plasma membrane and Golgi pools of β1-adrenergic receptors regulate different molecular and physiological functions in cardiomyocytes

Given that Golgi-generated cAMP specifically regulates PLB phosphorylation and lusitropy, we hypothesized that hormone-mediated cAMP responses by Golgi-localized β1AR should regulate cAMP-mediated PLB phosphorylation. Moreover, we hypothesized that β1ARs at the plasma membrane control downstream effectors in the vicinity of the plasma membrane. β1ARs localize to both the plasma membrane and Golgi membranes of neonatal and adult cardiomyocytes, as detected by immunostaining using an antibody against β1AR (Extended Data Fig. 3a-c). We have previously confirmed the specificity of the antibody using two different siRNAs against β1AR^19^. We further confirmed this immunostaining result using adult cardiomyocytes derived from β1AR/β2AR double knockout mice. We did not detect any fluorescence signal in either the Golgi or the plasma membrane in β1AR/β2AR double knockout cardiomyocytes (Extended Data Fig. 3c). We then tested whether this antibody is capable of detecting low levels of β1AR in transfected cell lines. HeLa cells were transfected with doxycycline-inducible promotor (Tet-on) β1AR construct and were treated with 0.1 and 0.5 μg/ml doxycycline for 24hour post transfection. Low levels of β1AR expressing cells were detected using the β1AR antibody used here (Extended Data Fig. 3d), further confirming the specificity of this antibody.

We previously reported that both the plasma membrane and the Golgi pool of β1ARs can promote cAMP generation^9,19^. In healthy cardiomyocytes, β1AR signaling regulates cardiac responses through PKA-mediated phosphorylation of proteins such as TnI, RyR2, and PLB (Extended Data Fig. 4a)^10^. We used immunofluorescence imaging to visualize cellular localization patterns of phosphorylated forms of RyR2, TnI, and PLB upon 10 μM epinephrine in cardiomyocytes. While phosphorylated PLB localized near the perinuclear/Golgi membranes (Extended Data Fig. 4b and c), phosphorylated TnI and RyR2 did not colocalize with the Golgi marker and showed a plasmalemma (the outer plasma membrane regions in cardiomyocytes) and near premature t-tubule (an invaginated region of the plasma membrane that is near the SR junctional region) localization pattern, respectively (Extended Data Fig. 4d-g). Based on this distinct localization pattern, we hypothesized that β1AR-mediated cAMP likely regulates distinct PKA effectors in each membrane compartment’s vicinity.

To assess whether the plasma membrane and Golgi pools of β1AR regulate different PKA effectors, we pharmacologically blocked β2AR with 10 μM ICI118551 to specifically test the function of β1AR in cardiomyocytes. We took advantage of membrane-permeant and impermeant agonists of β1AR to compare the functions of plasma membrane and Golgi-localized β1AR in adult cardiomyocytes (Extended Data Fig. 4a). We have previously demonstrated that epinephrine, a membrane impermeant βAR agonist, requires a monoamine transporter, organic cation transporter 3 (OCT3), to reach the Golgi lumen and activate Golgi-localized β1AR^19^. OCT3 is expressed in cardiomyocytes (Extended Data Fig. 5b)^51^. Pharmacological inhibition of OCT3 inhibits epinephrine/norepinephrine-mediated Golgi-localized β1AR activation^9,19^. Importantly, OCT3 inhibition abolishes epinephrine-mediated phosphorylation of PLB but not β1AR-mediated phosphorylation of TnI and RyR2 (Fig. 3a-d). Unlike epinephrine, dobutamine, a membrane-permeable β1AR agonist, can activate Golgi-localized β1AR independently of OCT3 (Fig. 3a, last lane and Extended Data Fig. 5c-f). Additionally, cardiomyocytes derived from OCT3 knock-out mice that have similar expression levels of β1AR as wild type cardiomyocytes (Extended Data Fig. 5b) showed no PLB phosphorylation upon epinephrine stimulation but displayed an increase in TnI and RyR2 phosphorylation (Fig. 3e-h). In contrast, dobutamine caused phosphorylations of TnI, RyR2, and PLB in OCT3 knock-out cardiomyocytes (Extended Data Fig. 6a-d). Thus, our results indicate that plasma membrane-localized β1AR regulates TnI and RyR2 phosphorylations to control inotropy (contraction), whereas Golgi-localized β1AR regulates PLB phosphorylation to control lusitropy (relaxation).

**Figure 3.**
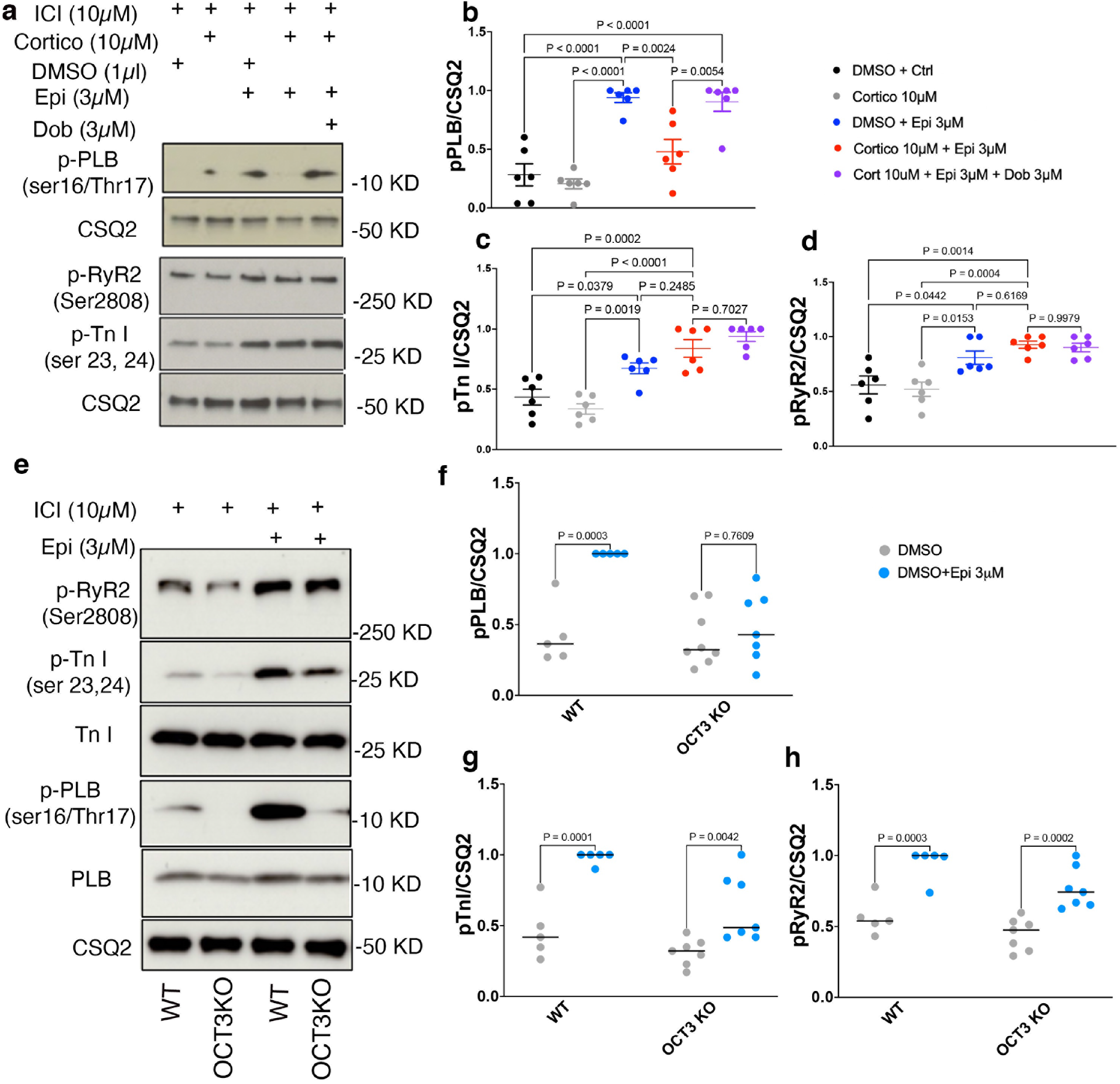
Plasma membrane and the Golgi pools of β1AR function differently in adult mouse cardiomyocyte. **a**. Corticosterone (Cortico)-mediated inhibition of OCT3, the transporter that allows epinephrine (Epi) to access the Golgi, blocks phosphorylation of PLB but not TnI and RyR2 in adult cardiomyocytes. Membrane permeable β1AR selective agonist, dobutamine (Dobut), promotes PLB phosphorylation independent of OCT3. Adult cardiomyocytes were pretreated with β2AR selective antagonist ICI-118551 (ICI) to isolate the function of β1ARs. Thus, Golgi β1AR regulates PLB phosphorylation, a mediator of cardiomyocyte relaxation. **b-d**. Quantification of immunoblots of p-RyR2 Ser2808, p-TnI Ser23/24, and p-PLB Ser16/Thr17 normalized to the protein level of CSQ2, and then reported as a percentage of the highest value in the groups. The quantified data from different experiments were presented as mean ± S.E.M. The p-values were calculated by one-way ANOVA. n=6 biological replicates. **e**. Representative western blots of phosphorylation profiles of RyR2, TnI, and PLB regulated by β1AR in adult cardiomyocytes derived from OCT3 (*SLC22A3*) knock-out mice and compared to wild-type. **f-h**. Quantification of immunoblots of p-RyR2 Ser2808, p-TnI Ser23/24, and p-PLB Ser16/Thr17 normalized to the protein level of CSQ2, and then reported as a percentage of the highest value in the groups. The quantified data from different experiments were presented as mean ± S.E.M. The p-values were calculated by two-way ANOVA. n=5 and 7 biological replicates for wild-type and OCT3 knock out cardiomyocytes, respectively.

Given that PLB is the key regulator of Ca^2+^ reuptake by SERCA channel, we then assessed Ca^2+^ dynamics in cardiomyocytes. We isolated adult cardiomyocytes and incubated them with Fluo 4 AM, a Ca^2+^ dye. Upon 1Hz field stimulation, both cardiomyocytes derived from the wild-type and OCT3 knock-out showed similar baseline Ca^2+^ dynamics (Extended Data Fig 7a and b). Epinephrine treatment increased the Ca^2+^ transient amplitude in both wild-type and OCT3KO-derived cardiomyocytes (Extended Data Fig. 7c). Wild-type adult cardiomyocytes also showed a decreased calcium decay tau in response to epinephrine stimulation, a readout of accelerated Ca^2+^ reuptake by SERCA upon PLB phosphorylation. However, adult cardiomyocytes isolated from OCT3 knock-out mice did not show a significant decrease in calcium decay tau upon epinephrine treatment (Extended Data Fig. 7d). These results are consistent with the lack of PLB phosphorylation observed in OCT3 knock-out cardiomyocytes upon epinephrine stimulation (Fig. 3e and f).

### β1AR autoantibodies specifically activate plasma membrane-localized β1AR and regulated cardiomyocytes contractility

To further distinguish the roles of Golgi and plasma membrane-β1AR signaling in regulating cardiomyocyte contractility, we took advantage of an autoantibody against β1AR to specifically activate β1ARs only at the plasma membrane. Autoantibodies against β1ARs have been reported in various cardiac diseases, including dilated cardiomyopathy^52–54^. Many of these autoantibodies function as agonists because their epitope sequences have sequence similarities to the extracellular loop 2 of β1ARs (Figure 4A)^54^. Previous studies have shown a measurable cAMP production and positive inotropic response upon treating cardiomyocytes with autoantibodies^55,56^. Given that antibodies are membrane impermeant and cannot cross the plasma membrane, we used them to activate the plasma membrane pool of β1AR specifically. We first tested whether this antibody functions as an agonist. To do this, we utilized a previously generated nanobody-based biosensor (Nb80-GFP) to detect active conformation of βARs^19,26^. Stimulating HEK293 cells expressing SNAP-tagged β1AR and Nb80-GFP with 100 nM antibody generated against extracellular loop 2 of β1AR resulted in subtle recruitment of Nb80-GFP to the plasma membrane-localized β1AR (Fig. 4a and Extended Data Fig. 8), suggesting that the β1AR antibody functions as a partial agonist. Additionally, we found a slight increase in cAMP concentration (~10 pmol/mg) when neonatal cardiomyocytes were treated with β1AR autoantibody, further supporting the notion that this antibody acts as a partial agonist (Fig. 4b). We then stimulated isolated cardiomyocytes with two different concentrations of the autoantibody (10 and 33 nM) and found an increased RyR2 phosphorylation but not PLB and TnI phosphorylation (Fig. 4c and d). These data further support a model where different pools of β1ARs regulate distinct functions in cardiomyocytes, with Golgi-localized-β1ARs regulating PLB phosphorylation and the plasma membrane/t-tubule-localized β1ARs regulating RyR2 phosphorylation. Interestingly, we did not observe TnI phosphorylation by β1ARs antibody at these concentrations. We speculate that this is due to lower levels of β1AR expression on the plasmalemma (the outer plasma membrane regions in cardiomyocytes) compared to t-tubule, an invaginated region of the plasma membrane that is near the SR junctional region. This is consistent with our immunostaining data showing distinct localization pattern of β1AR on the plasma membrane of neonatal cardiomyocytes in being concentrated in the premature t-tubules near the SR junctional region (dyadic cleft) where RyR2 is localized and very little staining on the plasmalemma (Extended Data Fig. 3b and c). Thus, the lack of TnI phosphorylation by the β1AR autoantibody suggests that low levels of cAMP generated by the activated β1AR on the premature t-tubule can only activate downstream PKA effector near the dyadic cleft (RyR2) and not PKA effectors that are outside of dyadic cleft (TnI and PLB).

**Figure 4.**
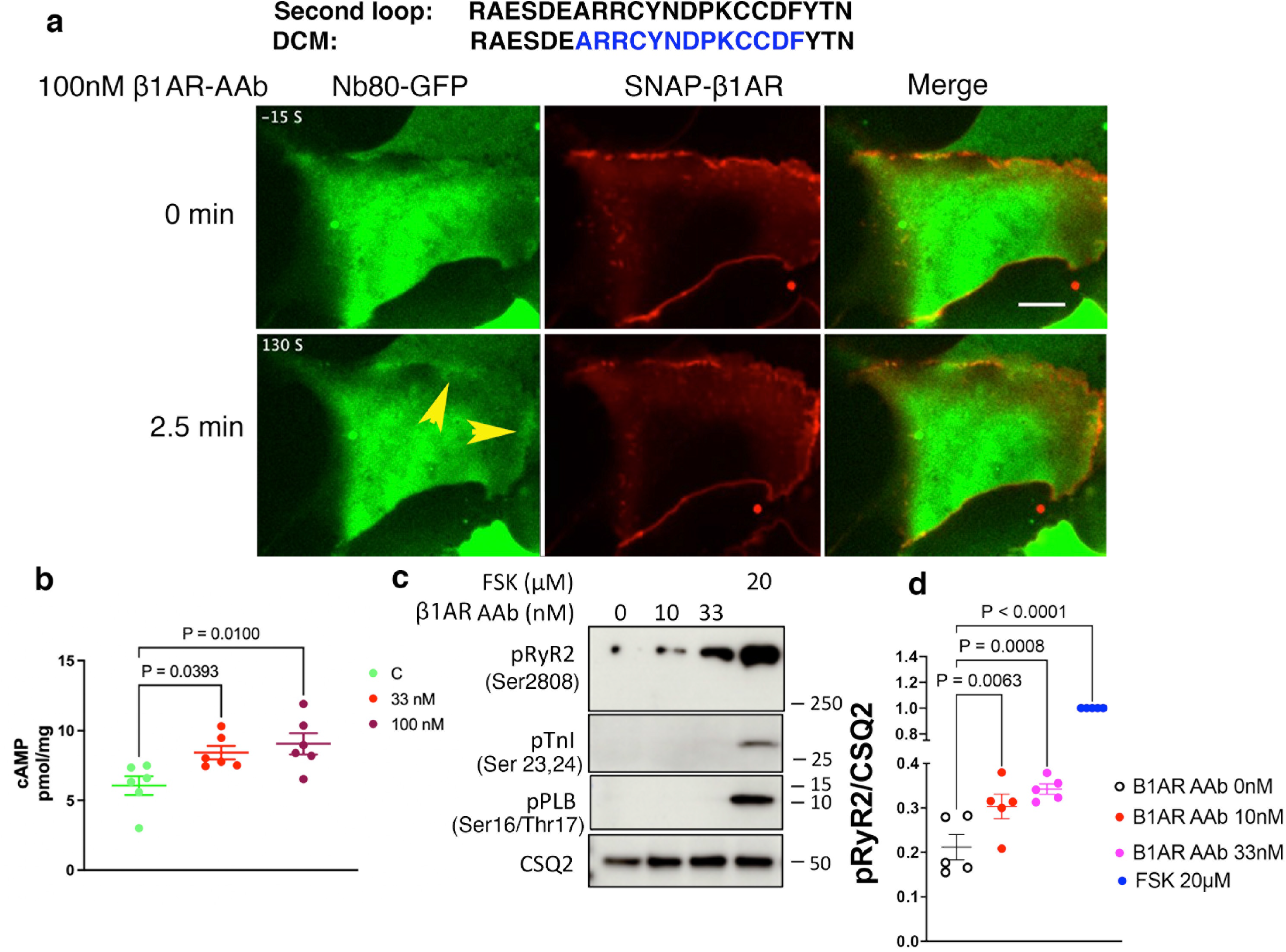
β1AR autoantibody specifically activate plasma membrane-localized β1AR and regulates RyR2 phosphorylation in neonatal mouse cardiomyocytes. **a**. The amino acid similarity between the second extracellular loop of β1AR and an autoantibody (AAb) epitope region (blue color labeled) found in patients with dilated cardiomyopathy (DCM) (top), representative images of HEK293 cells transfected with SNAP-tagged β1AR and nanobody based biosensor for βARs (Nb80-GFP) before and after 100 nM autoantibody stimulation (bottom). Nb80-GFP is recruited to the plasma membrane-localized β1AR after 2.5 minutes (arrowheads). Scare bar=10μm. **b**. cAMP generation mediated upon 33 and 100 nM autoantibody treatment in neonatal cardiomyocytes. Cells were stimulated for 15 minutes and lysed for direct cAMP determination by ELISA. cAMP concentrations were normalized to the relative protein concentrations in cell lysate of each sample. The quantified data are represented as mean ± S.E.M. The p-values were calculated by one-way ANOVA; C, control group. n=6 biological replicates **c**. Representative western blots of RyR2, TnI, and PLB phosphorylation profiles regulated by β1AR AAb in mouse neonatal cardiomyocytes. Mouse neonatal cardiomyocytes were treated with 10 and 33 nM β1AR AAb or 20μM FSK for 15 minutes. **d**. The protein levels of p-RyR2 Ser2808, p-TnI Ser23/24, and p-PLB Ser16/Thr17 were analyzed. The protein level of p-RyR2 was normalized with the protein level of CSQ2, and then reported as a percentage of the highest value in the FSK-treated group (20 *μ*M). The quantified data from different experiments were presented as mean ± S.E.M. The p-values were calculated by one-way ANOVA. n=5 biological replicates.

### Plasma membrane and Golgi PKAs have distinct functions in cardiomyocytes

PKA is a holoenzyme composed of two regulatory (PKA-R) and two catalytic (PKA-C) subunits anchored to membranes by A-kinase anchoring proteins. As a result, PKA holoenzymes are highly compartmentalized^57,35,41,58^. It was commonly believed that the PKA-C subunit dissociates from the PKA-R subunit in the presence of excess cAMP, and thus PKA-C can activate downstream effectors localized within the cells^59,60^. However, recent studies have revealed that the activity of the PKA-C subunit is constrained to targets within a radius of 15-25 nm^40^. Given that this spatially and functionally restricted PKA can only phosphorylate proximal downstream targets in cardiomyocytes, it stands to reason that the plasma membrane-localized receptors are unlikely to be the sole source of cAMP-mediated PKA activation. To test which pool of PKA within the cells regulates the phosphorylation of downstream effectors, we targeted a dominant-negative PKA (dnPKA), a constitutively repressive version of PKA-RIa that is insensitive to cAMP^61,62^ to the plasma membrane (PM-dnPKA) (Fig. 5a). To target dnPKA to the plasma membrane, we fused it to the CAAX motif of the K-Ras protein. PM-dnPKA was mainly localized on thin parallel striation that colocalized with sarcomeric z-disk markers (α-actinin), a membrane region in cardiomyocytes that closely coincides with t-tubules (Fig. 5b, arrowhead)^63–67^. We could also observe a fraction of PM-dnPKA on the plasmalemma of the plasma membrane in neonatal cardiomyocytes (Fig. 5b, arrow). We then assessed how the inhibition of the plasma membrane pool of PKA affects epinephrine-mediated phosphorylation of downstream PKA targets in cardiomyocytes. Stimulation of neonatal cardiomyocytes with epinephrine resulted in the phosphorylation of RyR2, TnI, and PLB. Interestingly, epinephrine-stimulated cardiomyocytes expressing PM-dnPKA showed abrogated RyR2 phosphorylation, but TnI and PLB phosphorylations remained unchanged (Fig. 5c-f). This is consistent with the observation that PM-dnPKA is largely concentrated on the pre-mature t-tubules near the localization of RyR2 but present at low levels in plasmalemma which is near to where TnI is localized (Fig. 5b). These data along with our findings using autoantibodies (Fig. 4c) further confirm that the plasma membrane-β1AR, most likely concentrated in the premature t-tubules, is distinctly activating PKA at the t-tubule, which then regulates RyR2 near the SR junctional regions.

**Figure 5.**
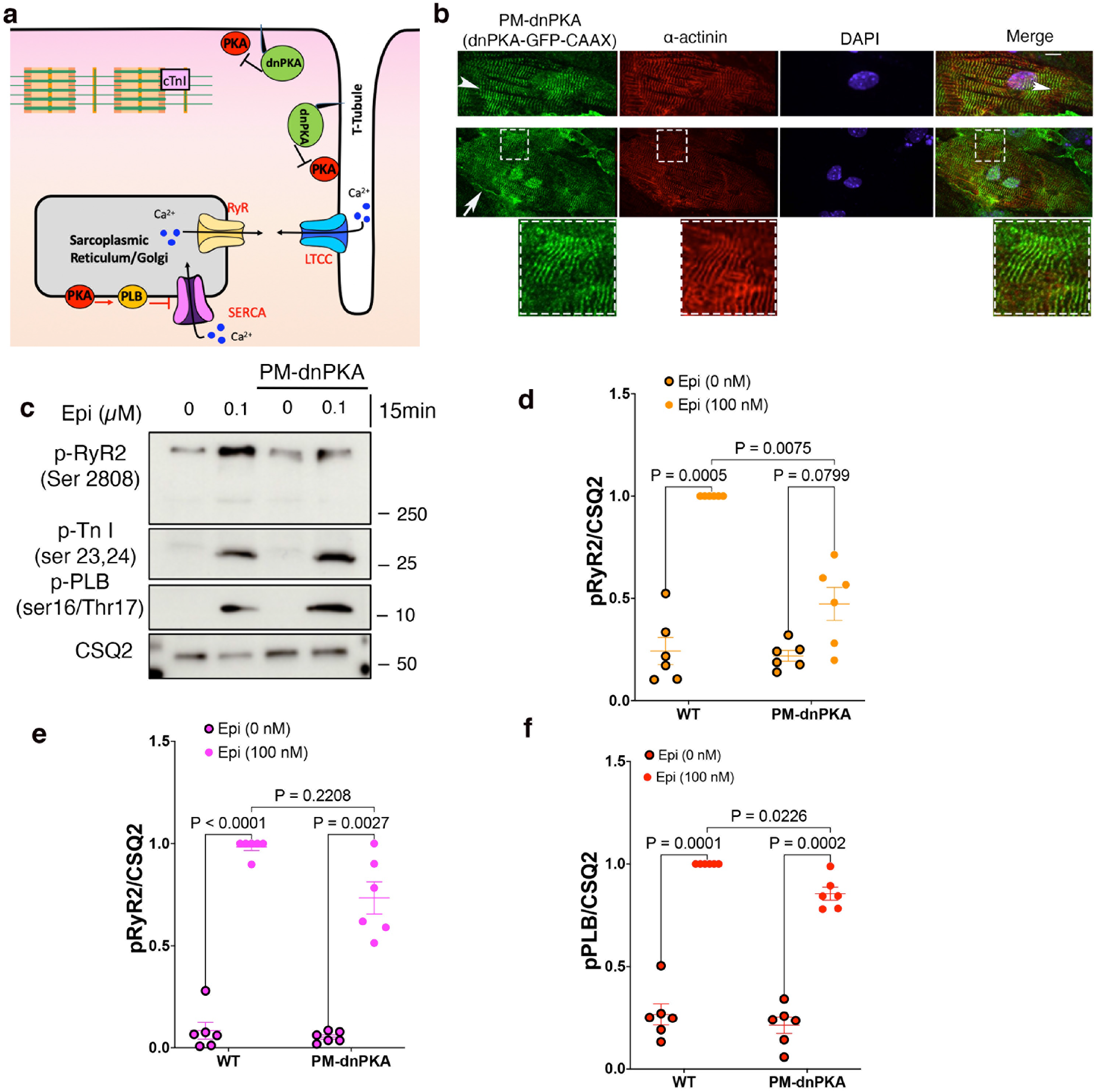
Plasma membrane and the Golgi pool of PKA have distinct functions. **a**. Model of targeting a dominant-negative PKA (dnPKA) to the plasma membrane (PM-dnPKA) to locally regulate PKA activity in cardiomyocytes. PM-dnPKA was generated by fusing dnPKA with sfGFP and CAAX motif. **b**. PM-dnPKA, visualized by GFP (green), was co-stained with α-actinin (red), a marker of t-tubules, and DAPI. Insets show co-localization of PM-dnPKA to t-tubules. Scale bar, 10 *μ*m. **c**. The representative Western blots of RyR2, TnI, and PLB phosphorylation profiles regulated by epinephrine in the absence or presence of PM-dnPKA expression. The protein levels of p-RyR2 Ser2808, p-TnI Ser23/24, and p-PLB Ser16/Thr17 were analyzed in wild-type and PM-dnPKA-expressing moue neonatal cardiomyocytes without or with 0.1 *μ*M epinephrine treatment for 15 minutes. **d-f**. The band intensities of p-RyR2, p-TnI, and p-PLB were normalized with CSQ2 intensity and then reported as a percentage of the highest value in the groups. The quantified data from different experiments were presented as mean ± S.E.M. The p-values were calculated by two-way ANOVA. n=6 biological replicates.

### OCT3 knock out mice have preserved inotropy but delayed lusitropy

Increased sympathetic activity during the fight and flight response or exercise causes an increase in epinephrine/norepinephrine levels in the circulation and enhances βARs activity. Thus, the heart efficiently augments cardiac output by increasing the heart rate, dromotropy (conduction speed), inotropy (force of contraction), and lusitropy (rate of relaxation). Our data in isolated adult and neonatal cardiomyocytes suggest that plasma membrane-localized β1AR regulates RyR2 phosphorylation, a key Ca^2+^ channel that increases the release of Ca^2+^ from SR, and thus triggers the cardiac muscle to contract. In contrast, we found that Golgi-localized β1AR specifically regulates PLB phosphorylation, a key regulator of Ca^2+^ reuptake to the SR through SERCA Ca^2+^ channels, thus promoting cardiac muscle relaxation (Fig. 3–5). These data suggest that plasma membrane β1AR regulates inotropy whereas Golgi-localized β1AR regulates lusitropy. To test this hypothesis, we performed real-time measurements of pressure and volume (PV loop) within the left ventricle of the mice in response to bolus injections of epinephrine (Figure 6a). Several physiologically relevant hemodynamic parameters, such as stroke volume, ejection fraction, myocardial contractility, and lusitropy, can be determined from these loops (Fig. 6a). To measure the PV loop upon stimulation of βARs, we inserted a 1.4-F pressure-conductance catheter and injected mice at the right jugular vein with 10 μg/kg epinephrine. To isolate the function of the plasma membrane and Golgi-localized β1AR, we compared wild-type and OCT3 knock-out mice (Extended Fig. 5a and b) ^68^. Given that OCT3 facilitates the transport of epinephrine to the Golgi-localized β1AR and OCT3 inhibition leads to abrogated PLB phosphorylation in cardiomyocytes (Fig. 3e-h), we predicted that OCT3 knock-out mice will have delayed lusitropic response. Epinephrine injection induced an increase in the heart rate of both wild-type and OCT3 knock-out mice (Supplementary Table 1). Importantly, the maximal rate of left ventricle pressure change (dp/dt max), ejection fraction, and cardiac output, which are the key indications of systolic function of contraction (inotropy), were similar upon epinephrine injection between the wild-type and OCT3 knock-out mice (Fig. 6b, c and f, Supplementary Table 1). An increase in contractility is observed as an increase in dP/dt max during isovolumic contraction. Thus, these data suggest that wild-type and OCT3 knock-out mice have a similar rate of contraction upon epinephrine injection. However, the minimal rate of left ventricle pressure change (dp/dt min), which is manifested as an increase in diastolic function or an increase in the rate of relaxation (lusitropy), was delayed in OCT3 knock-out mice (Fig. 6c and g, Supplementary Table 1). Moreover, tau, which represents the exponential decay of the ventricular pressure during isovolumic relaxation, was also delayed in OCT3 knock-out mice compared to wild-type (Fig. 6c and h, Supplementary Table 1). Importantly, injection of dobutamine, a membrane-permeant β1AR agonist that does not require OCT3 to activate Golgi-localized β1AR^9,19^, caused similar increases in the rate of contraction, relaxation, and tau in both wild-type and OCT3 knock-out mice (Fig. 6d-h, Supplementary Table 1). Altogether, these data suggest that OCT3 knock-out mice have preserved systole (inotropy) but delayed diastole (lusitropy) upon epinephrine stimulation.

**Figure 6.**
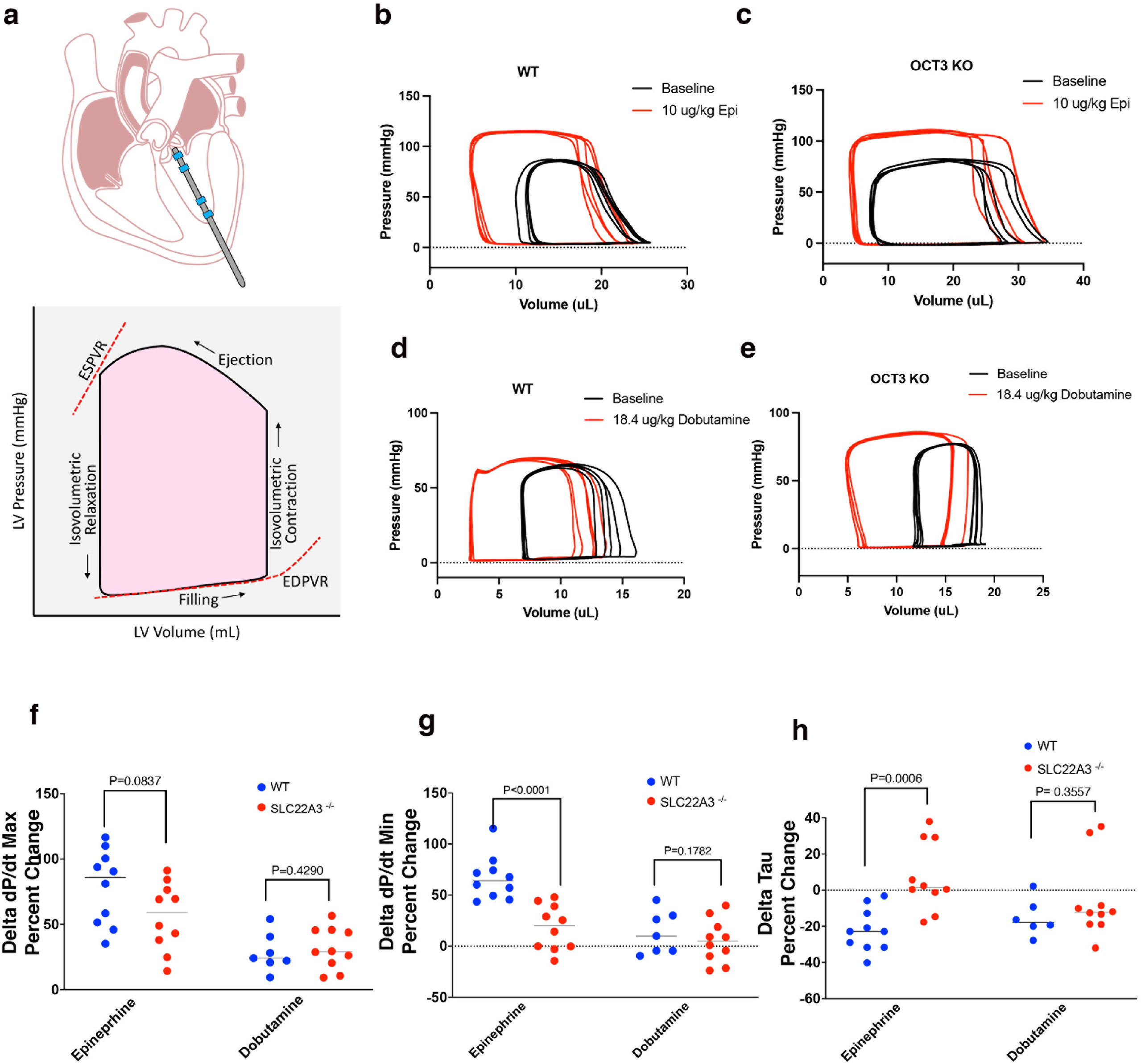
Pressure-Volume measurement of OCT3 knock-out mice revealed preserved systole but impaired diastole upon epinephrine stimulation. **a**. Diagram demonstrating the placement of the catheter for pressure-volume measurements in mouse hearts. The catheter (Millar Instruments) is inserted into the apex through a needle stab wound. The catheter has one pressure sensor and conductance electrodes which measure ventricular volume (top). Representative pressure-volume loop. Each loop shows the changes in pressure and volume during isovolumetric contraction, ejection, isovolumetric relaxation, and filling of the left ventricle. Shifts in the loops between basal and stimulated conditions can provide a comprehensive analysis of cardiac function and can be used to assess the heart’s performance by quantitatively measuring hemodynamic parameters. Multiple cardiac indicators, including the end-systolic pressure-volume relationship (ESPVR), end-diastolic pressure-volume relations (EDPVR), stroke volume, cardiac output, and ejection fraction, can be derived from PV loops (bottom). Representative hemodynamic pressure–volume loops (5 loops) upon 10μg/kg epinephrine injection in **b**. wild-type and **c**. OCT3 knock-out mice. Representative hemodynamic pressure–volume loops (5 loops) upon 18.4μg/kg dobutamine injection in **d**. wild-type and **e**. OCT3 knock-out mice. **f**. Pressure volume (PV) loop experiments were performed on mice to determine the effect of Golgi-delimited signaling of the β1AR. We examine this using OCT3 (*SLC22A3*) knock-out mice to limit the access of epinephrine (cell-impermeable endogenous β1AR agonist) to the Golgi compared with dobutamine (cell-permeable selective β1AR agonist) treated mice. Wild-type and OCT3 (*SLC22A3*) knock-out mice were anesthetized, and an apical catheterization of the left ventricle was performed. Basal PV-loops were collected as a control for each mouse. Either epinephrine (10 *μ*g/kg) or dobutamine (18.4 *μ*g/kg) were administered by bolus injection through the jugular vein and repeated measurements of PV loops were collected. The maximum dP/dt is the derivative of pressure that increases over time and is a measurement of systolic function. **g**. The minimum dP/dt is the derivative of pressure that decreases over time and is an indicator of relaxation of the left ventricle. **h**. Tau represents the decay of pressure during isovolumetric relaxation that is preload independent. Data are presented as mean ± S.D. The p values were calculated by t-test. n=10 wildtype and OCT3 knock-out mice for epinephrine treatment and n=7 wildtype and 10 OCT3 knock-out mice for dobutamine treatment.

## Discussion

Our findings demonstrate the cellular and physiological significance of cAMP generation at specific locales. We present evidence that cells with more complex architecture, such as cardiomyocytes, distinguish local cAMP generation and elicit different physiological outputs. We demonstrated that localized activation of β1ARs at subcellular compartments leads to local generation of cAMP and activation of downstream PKA effectors that are in the vicinity of each compartment. We found that cAMP generation at the Golgi results in PLB phosphorylation and consequently increased rate of relaxation in cardiomyocytes, an observation verified in the intact hearts of zebrafish. Furthermore, we found that the plasma membrane pool of cAMP regulates local PKA effectors, such as RyR2, leading to increased contractile force. Importantly, we showed that a monoamine transporter (OCT3) that facilitates the transport of epinephrine/norepinephrine regulates the activation of Golgi-β1AR-mediated PLB phosphorylation. Thus, epinephrine stimulation in OCT3 knock-out cardiomyocytes only activates the β1AR pools at the plasma membrane. This observation was further verified in OCT3 knock-out mice, where the force of contraction (systole) was preserved upon epinephrine injection, but the relaxation rate (diastole) was impaired.

Examining the publicly available phenotype across >420,000 individuals in UK Biobank with exome sequencing data shows that loss of function OCT3 (SLC22A3) is significantly associated with cardiovascular diseases, specifically diastolic pressure^69^. Recently, it was shown that several prescription drugs that potently inhibit OCT3 cause adverse reactions related to cardiovascular traits^70^. The findings from this study suggest the possibility that these adverse reactions may be due to alterations in subcellular cAMP/PKA signaling caused by inhibition of OCT3.

Our data using an antibody against the extracellular loop 2 of β1AR to specifically activate the plasma membrane-pool of β1AR supports the notion that β1AR is mainly concentrated at premature t-tubule near the SR junction region in neonatal cardiomyocytes where RyR2 is localized. Thus, generating cAMP by plasma membrane-pool of β1AR resulted in the phosphorylation of RyR2 and had no effect on PLB and TnI phosphorylation found in the non-junctional SR regions and myofilaments, respectively. Accordingly, inhibiting PKA at the premature t-tubules in neonatal cardiomyocytes, abrogates epinephrine-mediated RyR2 phosphorylation but had no significant effect on TnI and PLB phosphorylation. These findings suggest that the functional pool of β1AR/cAMP/PKA resides in the t-tubules, near the SR junctional region, thus regulating the increase in the contractile force of cardiomyocytes. The β1AR autoantibodies are present in more than 30% of patients with dilated cardiomyopathy. It has been reported that these autoantibodies function as an agonist and specifically induce a positive inotropic effect in isolated cardiomyocytes^56^. Our findings provide a potential mechanism for this observation.

There are several different genetically encoded membrane-localized fluorescence and bioluminescence-based biosensors that have been developed to study cAMP compartmentalization^71–74^. Almost all these studies have focused on the role of PDEs, AKAPs, and PKA in forming cAMP domains at different subcellular compartments but assumed that the sole source of cAMP are GPCRs that are activated on the plasma membrane. Previous views of localized-GPCR signaling have been mainly attributed to receptor-associated cAMP micro- or nanodomain localization on the plasma membrane. For instance, it has been shown that β2ARs, but not β1ARs, are exclusively associated with caveolae and lipid rafts^75–77^. Thus, it was thought that the distinct signaling functions of β1AR and β2AR are due to their unique localization on the plasma membrane^75,78–87^. More recently, the cAMP nanodomain formation on the plasma membrane has been reported for glucagon-like 1 peptide receptor and β2AR where signaling specificity is determined based on the formation of receptor-associated cAMP nanodomains on the plasma membrane ^38^. The significance of GPCR signaling at subcellular locations other than the plasma membrane has only recently been explored^27,88^. Nash *et al* have demonstrated that inhibition of OCT3 abrogates β1AR-mediated Epac-dependent phospholipase Cε activation and hydrolysis of phosphatidylinositol-4-phosphate, a signaling pathway that contributes to the hypertrophic responses. More recently, it has been reported that a pool of β1AR is associated with the SERCA2 complex and regulates calcium transients and contraction responses^29^. Our data here provide evidence for the physiological significance of cAMP nanodomain formation by activated GPCRs on the plasma membrane and the Golgi membranes for regulating distinct cardiac function.

In patients with heart failure, lusitropic effects of catecholamines appear to be exerted by lower concentrations than inotropic effects^89^. It is well established that cAMP compartmentation is disrupted in failing hearts, due to mis-localization of PKA and their corresponding AKAPs that tether PKA to discrete subcellular sites^90–94^. Whether this is due to reduced or enhanced activity of βARs subtypes at specific membrane locations is not known and requires further investigation^89^.

Our novel findings on the significance of local cAMP signaling generation by activated β1AR could have important implications for a better understanding of cardiac diseases. For instance, our PV-loop measurements of OCT3 knock-out mice upon epinephrine stimulation mimic what is seen in diastolic dysfunction, a highly significant but poorly understood clinical condition where the cardiac muscle contraction is preserved, but relaxation is impaired. Our findings raise the possibility that patients with preserved systole and impaired diastole could have aberrations in cAMP signaling from the Golgi caused by a reduced receptor pool at the Golgi, impaired expression or reduced plasma membrane localization of OCT3, or reduced activity of downstream PKA effectors such as PLB. Establishing the physiological significance of GPCR/cAMP signaling from subcellular compartments in healthy cardiomyocytes is the first step in unraveling how this signaling specificity goes awry to cause cardiac disease.

## Figure Legends

**Extended Data Figure 1. Golgi-bPAC stimulates cAMP generation in response to blue stimulation over time. a**. Representative images of HeLa cells expressing Golgi-bPAC. **b**. Phosphorylation profiles of RyR2 and PLB induced by Golgi-bPAC in mouse neonatal cardiomyocytes. The protein levels of p-RyR2 Ser2808 and p-PLB Ser16/Thr17 were analyzed in the Golgi-bPAC-expressing mouse neonatal cardiomyocytes kept in the dark or exposed to 0.34 or 3.20 *μ*W/cm^2^ blue light for 1.5, 3 and 7minutes. The protein level of Golgi-bPAC was analyzed using the SNAP antibody. The protein level of CSQ2 was used as a loading control. **c**. Dose-response curve of cAMP generation in mouse neonatal cardiomyocytes upon various epinephrine concentrations n=4 biological replicates. **d**. The subcellular localization of p-PLB before blue light illumination (top); the subcellular localization of p-PLB p-RyR2, p-TnI, upon stimulation of Golgi-bPAC with 3.20 *μ*W/cm^2^ blue light in mouse neonatal cardiomyocytes (bottom rows). The protein localizations of p-RyR2 Ser2808, p-TnI Ser23/24, total PLB, and pPLB Ser16/Thr17 antibody were visualized (green) with Golgi marker stained by GM130 (red). Scale bar, 10 *μ*m. **e**. Phosphorylation profiles of RyR2, TnI and PLB induced by Golgi-bPAC before and after treatment of 20μM PKA inhibitor (H89) in mouse neonatal cardiomyocytes. The phospho protein levels were detected using of p-RyR2 Ser2808, p-TnI Ser23/24 and p-PLB Ser16/Thr17 antibodies. The protein level of Golgi-bPAC was analyzed using the SNAP antibody. The protein level of CSQ2 was used as a loading control.

**Extended Data Figure 2. Contraction and relaxation responses of Control Tg(Flk:Ras-cherry)s896 zebrafish to blue light stimulation. a**. Representative agarose gel electrophoresis of wild-type (WT) and GalT-bPAC zebrafish genotyping. The plasmid for the Golgi-bPAC was cloned by fusing GalT, bPAC, myc, and m-cherry into a pminiTol2 plasmid. The plasmid was linearized and co-injected with transposase mRNA into the cell of a ZF embryo at the one-cell stage. DNA fragments of GalT, bPAC, and mApple, were amplified from caudal fin samples to sort the transgenic fish. DRER is the DNA extraction and genotyping control for ZF samples. **b**. Golgi-bPAC zebrafish (72hpf) generate cAMP in response to 3 minutes blue light (4.2 *μ*W/cm^2^). Zebrafish were lysed in 120 *μ*L of 0.1M HCl, and then cAMP was detected by a direct determination ELISA assay. 100 *μ*M FSK treated zebrafish was used as a positive control. The quantified data from different experiments are presented as mean ± S.E.M. The p-values were calculated by one-way ANOVA. n=4 biological replicates. **c**. Heart rate measurements of Golgi-bPAC zebrafish in response to 1, 3, and 5 minutes blue light (4.2 *μ*W/cm^2^). Heart rates were calculated by measuring the distance between two consecutive highest points of the peaks of each cycle. The quantified data from different experiments are presented as mean ± S.E.M. The p-values were calculated by one-way ANOVA. n=38 zebrafish **d**. Control (Tg(Flk:Ras-cherry)) zebrafish were stimulated with 4.2 *μ*W/cm^2^ blue light for 1, 3, and 5 min and changes in time of contraction and relaxation were calculated. The quantified data are represented as mean ± S.E.M., p-values were calculated by t-test. n=7 control Flk:Rash-cherry zebrafish. **e**. Golgi-bPAC zebrafish were pretreated with IBMX for 90 minutes and then exposed to 4.2 *μ*W/cm^2^ blue light at each time points. The quantified data are represented as mean ± S.E.M., p-values were calculated by t-test. n=17.

**Extended Data Figure 3. Plasma membrane and Golgi localization of β1AR in mouse neonatal and adult cardiomyocytes. a**. Representative images of β1AR (green) localization relative to TGN38 (trans-Golgi) and GM130 (cis-Golgi) in mouse neonatal cardiomyocytes. **b**. Representative images of β1AR (green) localization relative to α-actinin (z-disk marker) in mouse neonatal cardiomyocytes. Endogenous β1AR is expressed on the plasma membrane/premature t-tubules and the Golgi membranes. **c**. Isolated adult cardiomyocytes from wild-type showed distinct t-tubule and Golgi localization, as detected by β1AR (green) and TGN38 (red) (Golgi marker) antibodies. The distinct t-tubules and Golgi staining of β1AR are not detectable in adult cardiomyocytes isolated from β1AR/ β2AR double knock-out mice. **d**. Representative images of HeLa cells transfected with a doxycycline-inducible promotor (Tet-on) controlled β1AR construct, treated with 0.1 and 0.5 μg/ml doxycycline for 24 hours post transfection. β1AR (green) was detected using β1AR antibody. Scale bar, 10 *μ*m.

**Extended Data Figure 4. Epinephrine stimulation of mouse neonatal cardiomyocytes reveals spatially distinct phosphorylation patterns of downstream PKA effectors. a**. Model of compartmentalized β1AR’s regulation of PKA effectors. The organic cation transporter 3 (OCT3) facilitates the uptake of epinephrine/norepinephrine to the Golgi-localized β1AR. **b-g**. Representative images of epinephrine-mediated phosphorylation of PLB, TnI, and RyR2 in neonatal cardiomyocytes. Neonatal cardiomyocytes were incubated with epinephrine (10 μM) for 20 minutes and immune-stained for p-RyR2 Ser2808, p-TnI Ser23/24, or p-PLB Ser16/Thr17, and TGN38. Representative ROIs in the merged images were analyzed by fluorescence line scan intensity and shown in the corresponding graphs (D, F, and H). The maximal fluorescence intensity of the PLB, TnI, and RyR2, relative to the Golgi markers, are measured along the width of the neonatal cardiomyocytes. The length of each ROI was normalized and organized into 100 bins; the average intensity of each bin is shown. These graphs demonstrate the phosphorylated proteins’ localization, spread, and intensity throughout the cells. n=15 cells per condition. Scale bar, 10 *μ*m.

**Extended Data Figure 5. Membrane permeable β1AR selective agonist, dobutamine, promotes PLB phosphorylation independent of OCT3 inhibition by corticosterone. a**. Representative agarose gel electrophoresis for genotyping of wild-type, heterozygous, and homologous OCT3 (*SLC22A3*) knock-out mice. **b**. Western blot of OCT3 (*SLC22A3*) (top) and β1AR (bottom) and CSQ2 from wild-type and OCT3 (*SLC22A3*) knock-out-derived cardiomyocytes. **c**. Representative western blots of adult cardiomyocytes pretreated with 10 *μ*M corticosterone (cortico) and 10 *μ*M ICI-118551 (ICI), then stimulated with 3 μM dobutamine (Dob). **d-f**. Quantification of Western blot of p-RyR2 Ser2808, p-TnI Ser23/24, and p-PLB Ser16/Thr17 normalized to the protein level of CSQ2, and then reported as a percentage of the highest value in the groups. The quantified data from different experiments were presented as mean ± S.E.M. The p-values were calculated by two-way ANOVA. n=6 biological replicates.

**Extended Data Figure 6. Membrane permeable β1AR selective agonist, dobutamine, promotes PLB phosphorylation in adult OCT3 knock-out cardiomyocytes. a**. Representative western blots of adult cardiomyocytes derived from wild-type or OCT3 knock-out stimulated with 0.3 or 3 μM dobutamine (Dob). **b-d**. Quantification of Western blot of p-RyR2 Ser2808, p-TnI Ser23/24, and p-PLB Ser16/Thr17 normalized to the protein level of CSQ2, and then reported as a percentage of the highest value in the groups. The quantified data from different experiments were presented as mean ± S.E.M. The p-values were calculated by t test. n=6 biological replicates.

**Extended Data Figure 7. Epinephrine-mediated Ca**^**2+**^ **decay tau was delayed in adult cardiomyocytes derived from OCT3 knock out mice. a**. Adult cardiomyocytes were loaded with 5 μM Ca^2+^ indicator (Fluo-4 AM). Representative line scan images and the corresponding trace of fluorescence obtain from epinephrine-mediated Ca^2+^ response upon 1Hz field stimulation in adult cardiomyocytes derived from wild-type (left) and OCT3 knock-out mice (right). **b-c**. Ca^2+^ transient amplitude, and **d**. decay tau were recorded with 1Hz pacing and upon 1 μM epinephrine stimulation for 5 minutes in adult cardiomyocytes derived from wild-type and OCT3 knock-out mice. The quantified data from different experiments were presented as mean ± S.E.M. The p-values were calculated by two-way ANOVA. n=6 and 7 cells for amplitude and decay tau, respectively, 3 biological replicates.

**Extended Data Figure 8. Nb80-GFP is recruited to activated β1AR on the plasma membrane upon autoantibody stimulation**. Representative images of HEK293 cells transfected with SNAP-tagged β1AR and nanobody based biosensor for βARs (Nb80-GFP) upon 100 nM autoantibody stimulation (bottom). Nb80-GFP is recruited to the plasma membrane-localized β1AR after 30 minutes (arrowheads). Scare bar, 10μm.

**Supplementary Table 1. Summary of raw data collected for pressure-volume measurement of wild-type and OCT3 knock-out mice**. Heart rate, maximal and minimal rate of left ventricle pressure change (dp/dt Max and dp/dt Min), tau, ejection fraction, and cardiac output of both wild-type and OCT3 knock-out mice were measured upon injection of epinephrine and dobutamine.

## Material and Methods

### Reagents and antibodies

Human insulin, human transferrin, and sodium selenite (ITS), urethane, 2,3-Butanedione monoxime (BDM), Taurine, protease XIV, polybrene, forskolin, epinephrine, dobutamine, corticosterone, and IBMX are from Sigma. Glutamax solution, Penicillin and Streptomycin, HEPES buffer, HBSS buffer, M199 medium, ultrapure H_2_O, Dulbecco’s minimal essential medium (DMEM), mouse laminin, and Halt™ protease and phosphatase inhibitor cocktail are from Thermo Fisher Scientific. Fetal bovine serum (FBS) and Nu serum IV are from Corning. Glucose, sodium chloride, potassium chloride, sodium phosphate monobasic monohydrate, magnesium chloride hexahydrate, Tris-base, K-pipes, HEPES, EDTA, DTT, DMSO, and Tween-20 are from Fisher Bioreagents. Calcium chloride, Trolox, and tricaine are from Acros Organics. ICI-118551 are from TOCRIS. Doxycycline is from Takara. Heparin solution is from Fresenius Kabi. Collagenase II is from Worthington. Bovine serum albumin (BSA) and dry milk powder is from Research Product International. EGTA is from Alfa Aesar. Triton X-100 is from Bio-Rad. Proteinase K is from Roche. Rabbit anti-phospho phospholamban (Ser16/Thr17) antibody (#8496) and rabbit anti-phospho troponin I (Ser23/24) antibody (#4004) are from Cell Signaling. Rabbit anti-phospho ryanodine receptor 2 (Ser2808) antibody (#PA5-104444) is from Thermo Fisher Scientific. Rabbit anti-Calsequestrin 2 antibody (#18422-1-AP) and Rabbit anti-SLC22A1 antibody (#24617-1-AP) are from Proteintech. Rabbit anti-SLC22A3 antibody (#ab183071) and rabbit anti-β1AR (#ab3442) are from Abcam. Goat anti-β1AR antibody is from Everest Biotech (EB07133). Rabbit anti-SNAP tag antibody (#P9310S) is from New England BioLabs. Mouse anti-GM130 is from BD Biosciences (#610822). Sheep anti-TGN38 is from Bio-Rad (#AHP499G). Mouse anti-α-actinin is from Sigma (#A7811). Donkey anti-Mouse IgG (H+L) Highly Cross-Adsorbed Secondary Antibody, Alexa Fluor™ 647, (#A31571), Donkey anti-Rabbit IgG (H+L) Highly Cross-Adsorbed Secondary Antibody, Alexa Fluor™ 647 (#A31573), Donkey anti-Sheep IgG (H+L) Cross-Adsorbed Secondary Antibody, Alexa Fluor™ 488 (#A11015), Donkey anti-Rabbit IgG (H+L) Highly Cross-Adsorbed Secondary Antibody, Alexa Fluor™ 488 (#A21206), Donkey anti-Mouse IgG (H+L) Highly Cross-Adsorbed Secondary Antibody, Alexa Fluor™ 488 (#A21202) and Donkey anti-Sheep IgG (H+L) Cross-Adsorbed Secondary Antibody, Alexa Fluor™ 555 (#A21436) were purchased from Thermo Fisher Scientific.

### Plasmid construction

pLVXTetOne lentiviral vector (a gift from Dr. Jura lab) was used for doxycycline-induced protein expression in mouse neonatal cardiomyocytes. To generate pLVXTetOne signal peptide (SS)-SNAP-TGNP-bPAC plasmids, DNA fragments of SS-SNAP were amplified from pcDNA3_SS-SNAP-ADRB2 (a gift from Dr. von Zastrow lab). The fragments of bPAC and TGNP were amplified from cytoplasmic-bPAC (a gift from Dr. Reiter lab) and pmApple-TGNP-N-10 (Addgene plasmid #54954), respectively. To generate pLVXTetOne-dnPrkar1a-msfGFP-CAAX, the DNA fragments of dnPrkar1a, msfGFP, and CAAX were cloned from pCS2+dnPKA-GFP (a gift from Randall Moon, Addgene #16716), msfGFP containin plasmid (a gift from Dr. Giacomini lab), and pHR-SFFVp-CIB-GFPCAAX (a gift from Dr. Weiner lab). To generate pLVXTetOne_SS-SNAP-β1AR, SS-SNAP was cloned as previous description and β1AR were cloned from pcDNA3_SS-FLAG-β1AR (a gift from Dr. von Zastrow lab). The cloned DNA fragments were inserted into the pLVXTetOne lentiviral vector (a gift from Dr. Jura lab). To generate the pminiTol2 cmlc2: GalT-bPAC, the bPAC was amplified from cytosolic bPAC (a gift from Dr. Reiter lab), GalT and mApple were amplified from the FKBP-GalT-mApple plasmid^19^, and inserted into the pminiTol2 cmlc2 vector (a gift from Dr. Von Zastrow lab).The DNA fragments were amplified by Pfu Ultra II Hotstart PCR master mix (Agilent Technologies) and ligated with each respective vector by NEBbuilder HiFi DNA assembly master mix (New England BioLabs).

### Cell culture and lentivirus production

HEK293, HeLa, and HEK293T cells are cultured in DMEM (#11965092) containing 10% FBS. Lentiviral vector was co-transfected with pSPAX2 and pMD2.G plasmids (gifts from Dr. Julius lab) to HEK293T by TransIT-Lenti transfection reagent (Mirus Bio). The lentivirus was produced in DMEM containing 10% FBS and 1% BSA and then concentrated by the Lenti-X concentrator (Takara Bio).

### Animals

CD-1, wild-type C57BL/6, *Slc22a3*-null C57BL/6, and *Adrb1*^*tm1Bkk*^ *Adrb2*^*tm1Bkk*^/J mice (#003810) were housed in the facilities controlled by standardized environmental parameters, including a 12-hour light/dark cycle in 7 days per week, humidity 30-70%, temperature 20-26°C, and access to water and foods *ad libitum*. All animal experiments were approved by the Institutional of Animal Care and Use Committee of the University of California, San Francisco. Genotyping of wild-type and *Slc22a3* null alleles were performed as previous described^95^. The primer sets for the genotyping are: wild-type allele (F: 5’-gttctggcctaggcagtgcctctaat-3’ and R: 5’-gtgctaatgacaacacatggagatg-3’; 300bp) and Slc22a3-null allele (F: 5’-ggtactattcctcttgccaatcc-3’ and R: 5’-gtgctaatgacaacacatggagatg-3’; 500bp). Genotyping of *Adrb1*^*tm1Bkk*^ *Adrb2*^*tm1Bkk*^/J mice was performed based on the protocols and primer information on The Jackson Laboratory website. Genotyping was performed using GoTaq Green master mix (Promega).

Zebrafish were reared and handled in compliance with standard laboratory practices and IACUC protocols. Embryos were maintained in egg water at 28°C in the dark for 5 days, and then raised in a 14 h light/ 10 h dark cycle. Experimental embryos were assayed within 72hpf at which time sex cannot be easily identified. However, sex is unlikely to affect the signaling pathways and physiological outputs in this study. GalT-bPAC fish were generated through the Tol2 transposon transgenesis of an established zebrafish line, Tg(Flk:Ras-cherry)^s896 96,97^ Embryos were co-injected (PV pneumatic pico pump) at the one-cell stage with the pminiTol2 cmlc2: GalT-bPAC (4.5 pg) linear plasmid and capped transposase RNA (6.3 pg). Embryos positive for cherry fluorescence were sorted and genotyped. Genotyping to identify bPAC and Drer_Chr1 (DNA extraction control) were performed using GoTaq Green master mix (Promega). DNA samples for genotyping were extracted (lysis buffer: 10 mM Tris pH 8 2 mM EDTA 0.2% Triton X-100 200 μg/ml Proteinase K) from adult caudal fin clipping. The primer pairs used for genotyping are: bPAC (F: 5’-gtcaaccggtacttcagcatct-3’ and R: 5’-tcgtagtacttctgggcctcat-3’; 473bp), GalT (F: 5’-tgatccggcagaccctggaa-3; and R: 5’-gccctcgatctcgaactcgt-3’:470bp), mApple (F: 5’-ggctccaaggtctacattaagcac-3; and R: 5’-tgtagtcctcgttgtgggac-3’:424bp), Drer_ch1 (F: 5’-tatacgcggccataagtactga-3’ and R: 5’-gttcatttggggctttgggtat-3’; 218bp). To determine mating pairs of GalT-bPAC fish, cAMP measurements were performed. Embryos at 72 hpf obtained from each pair were incubated with IBMX (100 *μ*M, 90 min, 28°C). Anaesthetization by incubation of tricaine (0.04% w/v) was confirmed by a reflex test of the tail. Embryos were then exposed to 4.2 μW/cm^2^ blue light to stimulate GalT-bPAC or maintained in the dark for 5 minutes. cAMP detection was measured by a direct cAMP ELISA assay. Mating pairs that produced embryos that robustly generated cAMP in response to blue light were subsequently used in imaging experiments.

### Primary culture of cardiomyocytes

The processes for neonatal cardiomyocytes isolation are modified from previous research^98^. Briefly, hearts harvested from P1-2 neonatal CD1 pups were tear into small pieces in the ice-cold HBSS containing 20 mM HEPES. Heart pieces mixed with 225 IU/ml collagenase II were incubated on the tube rotator at 37°C for 5 minutes. After 10-time pipetting, the released cells in the buffer were collected by centrifuge at 500 xg for 5 minutes. The undigested heart tissues were digested again as describe above until the undigested tissue became white and the size do not decrease. The cells from each digestion were pooled together and resuspended in the neonatal cardiomyocyte culture media, which is DMEM (#11995065) containing 10% FBS, 10% Nu Serum IV, 10 mM HEPES, 10 mM Glutamax, Penicillin and Streptomycin and ITS. The released cells that pass through 40 *μ*m strainer plated on the regular dish to remove the most of fibroblasts at 37°C for 2 hrs. The suspended cells were collected and plated on the mouse laminin-coated dish. For the virus transduction, lentivirus was mixed with the culture media with polybrene (8 *μ*g/ml). The lentivirus was removed after 1-day transduction. The transduced neonatal cardiomyocytes were further treated with doxycycline for 3 days.

Adult cardiomyocytes were isolated from 2-3-month-old C57BL/6 wild-type and *Slc22a3* knockout mice using the Langedorff-free method^99^. The heparin solution was intraperitoneally injected into mouse (5 U/g). After 10 minutes, urethane, dissolved in 0.9% NaCl, was also intraperitoneally injected into mouse (2 mg/g). When the mouse was fully euthanized, the mouse heart was exposed, and the inferior vena cava was cut to release the blood. After the injection of EDTA buffer (130 mM NaCl, 5 mM KCl, 0.5 mM NaH_2_PO_4_-H_2_O, 10 mM HEPES, 10 mM Glucose, 10 mM BDM, 10 mM Taurine, in ultrapure H_2_O) into the right ventricle, the aorta was clamped. The clamped heart was moved to the EDTA buffer containing dish and then the EDTA buffer was injected into the left ventricle. Then, the clamped heart was moved to the perfusion buffer (130 mM NaCl, 5 mM KCl, 0.5 mM NaH_2_PO_4_-H_2_O, 10 mM HEPES, 10 mM Glucose, 10 mM BDM, 10 mM Taurine, 1 mM MgCl_2_-6H_2_O, in ultrapure H_2_O) containing dish and the perfusion buffer was injected into left ventricle. The clamped heart was further moved to the digestion buffer (perfusion buffer with 0.5 mg/ml Collagenase II and 0.05 mg/ml protease XIV) containing dish and then the digestion buffer was injected in to left ventricle. After digestion, the heart was tear into small pieces and gently triturated to dissociate the cardiomyocytes. The digestion processes were stopped by adding stop buffer (perfusion buffer with 5% FBS) and the suspended cardiomyocytes were pass through the 100 *μ*m strainer. The cardiomyocytes were enriched by gravity sedimentation and reintroduced calcium gradually. The cardiomyocytes were resuspended by plating media (M199 media with 5% FBS, 10 mM BDM, Penicillin and Streptomycin) and plated on the mouse laminin-coated wells at 37°C for 1 hours. After washing out the unattached cells by culture media (M199 media with 0.1% BSA, 10 mM BDM, Penicillin and Streptomycin, and ITS), the cardiomyocytes were cultured in culture media for further use.

### Blue light stimulation for activating bPAC protein in cardiomyocytes

After 1-day transduction, neonatal cardiomyocytes were treated with 100 ng/ml doxycycline for three days and then treated with 100 *μ*M Trolox for 4 hours. bPAC-expressing neonatal cardiomyocytes were put under the blue LED board in the incubator. After blue light stimulation for indicated interval, the neonatal cardiomyocytes were washed by ice-cold PBS once and lysed. To measure the blue light intensity, we used a Digital Handheld Optical Power and Energy Meter Console (#PM100D, Thorlabs) with a Slim Photodiode Power Sensor probe (S130C, Thorlabs). The light intensities were calculated from the power measured (W) and the probe detection surface of 0.7855cm2.

### Fixed-cell confocal imaging

For the staining of Golgi-bPAC and PM-dnPKA in the neonatal and adult cardiomyocytes or HeLa cells, cells were washed with PBS once and fixed with 3.7% formaldehyde in PEM buffer (80 mM K-PIPES pH 6.8, 1 mM MgCl_2_, and 1 mM EGTA) for 20 minutes at room temperature. For the staining of p-RyR2 Ser2808, p-TnI Ser23/24, pPLB Ser16/Thr17, PLB, and β1AR, cells were pre-permeabilized with 0.05% saponin diluted in PEM buffer on ice for 5 minutes before fixation. Fixation was performed using 3% paraformaldehyde diluted in PBS for 10 minutes at room temperature further quenched by 50 mM NH_4_Cl diluted in PBS for 10 minutes. Fixed cells were incubated with the primary antibody at room temperature for 1 hour or at 4°C for O/N in TBS containing 0.1% Triton. After the incubation with the secondary antibody at room temperature for 30 minutes, cells were mounted using antifade mounting medium with DAPI (Vector Laboratories). The images were taken by Nikon spinning disk confocal microscope.

### Lysate preparation, SDS-PAGE, and Western blot analysis

After the treatments, the cardiomyocytes from neonatal and adult mice were collected and lysed by RIPA buffer containing inhibitors of proteases and phosphatases at 4°C for 30 minutes on the tube rotator. Supernatants were collected after centrifuging at 4°C for 10 minutes and the protein amounts were determined by BCA assay (Sigma). The proteins were denatured by boiling for 10 minutes in the DTT containing sample buffer and separated by 4-20% Mini-PROTEIN TGX gels (BIO-RAD) and then transferred to the 0.2 *μ*m PVDF membrane (BIO-RAD). The PVDF membrane was further blocking by TBST (TBS buffer with 0.1% Tween-20) containing 3% milk at RT for 1 h and then incubated with the primary antibody in TBST containing 5% BSA at 4°C for O/N. The PVDF membrane was washed by TBST three times and then incubated with secondary antibody diluted in TBST containing 3% milk at RT for 1 hr. The unbonded secondary antibodies was removed by three times washing using TBST. The protein signals were visualized by ECL substrate (Thermo Fisher Scientific). To quantify band intensities, we first scan our films and convert them to an 8-bit format. We then inverted these images using imageJ software. The bands were selected using rectangular selection tool on ImageJ. The band intensities were measured, and each peak was separated by straight line selection tool. The area of each peak was measured by the Wand tool. Quantified phosphoprotein bands were then normalized to the total lysate bands (CSQ2) and data were presented as percentage of the maximum value that we have measured on each western blot.

### Intracellular calcium Imaging

Local increase in intracellular Ca^2+^ concentration was recorded using line-scanning confocal of isolated adult cardiomyocytes. Adult cardiomyocytes plated in Tyrode’s solution (140 mM NaCl, 4 mM KCl, 1.1 mM MgCl2, 10 mM HEPES, 10 mM glucose, 1.8 mM CaCl2; pH=7.4, with NaOH) and were loaded with 5 μM Ca^2+^ dye Fluo-4 acetomethyl ester (Fluo-4 AM) (Invitrogen Life Technologies) for 20 minutes at room temperature. Images were obtained using a Zeiss LSM 780-FLIM confocal microscope (40X water objective). Fluo-4AM was excited at 488 nm and emitted fluorescence was collected at 505 nm. The confocal pinhole is set to render 1.4 μm section. The gain was set at 800. To obtain intracellular Ca^2+^ transients, adult cardiomyocytes loaded with Fluo-4AM were electrically excited at 1 Hz for 15 s by filed stimulation using Ionoptix MyoPacel filed stimulator. A line scan image (256 pixel) were acquired at a rate of 0.473 ms per line along the longitudinal axis of the cells. The fluorescent values (F) were normalized to the basal fluorescence (F0) to obtain the fluorescence ratio F/F0 before and after 1 μM epinephrine stimulation. The Ca^2+^ decay tau was calculated by fitting the decay trace of 80 percent of the maximum response, using Graphpad Prism v9.3.1.

### Zebrafish imaging and analysis

Dechorionated embryos at 72 hpf were pretreated with a DMSO control (0.1% v/v) or IBMX (100 *μ*M, 90 minutes) and were anaesthetized with tricaine (0.04% w/v). Embryos were then mounted onto glass bottom imaging dishes (35 mm, MatTek) with low melting agarose (1% w/v) and maintained in tricaine for the duration of the experiment. All image series were acquired at 61 fps for 500 frames (λ_ex_= 561 nm, λ_em_=640 nm) with a Plan Apo 40X air objective (Nikon) on a spinning disk confocal (Nikon Eclipse Ti). Baseline images were obtained and GalT-bPAC was stimulated by exposing embryos to 4.2μW/cm^2^ of blue light for timepoints up to 5 minutes. The data were analyzed using FIJI v1.53f by measuring the fluorescence intensity of a ventricular ROI and determining the time between peak systole and peak diastole. Graphs were generated using Graphpad Prism v9.3.1. All data are expressed as Δ time (ms) relative to the baseline, as a mean ± S.E.M.

### Mouse cardiac pressure-volume loop acquisition and analysis

In order to assess ventricular systolic and diastolic function, we conducted pressure-volume loop experiments using a conductance catheter (Millar Instruments, Houston, TX)in mice as described in a previous study ^100^. Briefly, pressure and conductance calibrations were performed. Mice were initially anesthetized by inhalation of isoflurane (1.5% mixed with 100% oxygen). An endotracheal tube was placed and connected to the ventilator, and ventilator settings were based on animal weight ^101^. Mice were placed on a heating pad and body temperatures were maintained at 37ºC. Subsequently, analgesia was administered by subcutaneously injection of buprenorphine (0.05 mg/kg). Proper anesthetization was confirmed by a reflex test of the tail. 1 mg/kg Pancuronium (Sigma Life Science) was injected intraperitoneally to prevent respiratory artifacts during recordings ^102^. The aortic arch and inferior vena cava were exposed with a 6-0 silk ligature placed underneath separately. The right jugular vein was cannulated for subsequent fluid and medication infusion. A thoracotomy was performed, and the pericardium was bluntly dissected to expose the left ventricular apex. A 25-gauge needle was used to make a stab incision of the apex, followed by the insertion of a 1.4-F pressure-conductance catheter (PVR-1035, Millar Instruments, Houston, TX) through the incision. The intra-ventricular catheter position was optimized until rectangular-shaped loops were obtained (LabChart 8.5 Pro). Then, 200 *μ*L 0.9% saline was perfused slowly to replace body fluid loss. After steady state conditions were reached in 10 minutes, Baseline PV loops were recorded with three cycles of inferior vena cava and transverse aorta occlusion in sequence. Then epinephrine (10 *μ*g/kg) or dobutamine (18.4 *μ*g/kg) were injected and the above steps were repeated. To estimate G_p_, 10 *μ*L hypertonic saline (15% NaCl) were rapidly injected at the end of experiment. After 5 minutes, the blood was collected from the right ventricle for a cuvette calibration to transform conductance to volume. Cardiac parameters were obtained by offline data analysis on LabChart software (8.5).

### cAMP determination

Two modes of cAMP determination were performed. A direct cAMP ELISA kit (Enzo) was used in endpoint experiments and the pGloSensor (−20F) luminescence assay (Promega) was used for kinetic cAMP measurements. For time dependent cAMP production assays in HEK293 cells, Cells were transiently transfected with GalT-bPAC and the pGloSensor-20F plasmid. 24h after transfection, cells were incubated with Glosensor cAMP reagent for 1 h at 37ºC. For GalT experiments, cells were exposed to 4.2*μ*W/cm^2^ blue light for up to 300s. Luminescence measurements were acquired at 2 min intervals. Three baseline measurements were acquired after which cells were stimulated and measured for at least 20 min. Where specified cells were treated with, the PDE inhibitor IBMX (100 *μ*M), and positive control, forskolin (10 *μ*M). For primary cell and zebrafish cAMP determination ELISA assays were performed to determine cAMP. To measure epinephrine-induced cAMP production in mouse neonatal cardiomyocytes, cells were treated with a range of epinephrine (2 min, 37°C). Cells were immediately lysed in 175 *μ*L of 0.1M HCL lysis buffer for the direct cAMP determination by ELISA which was performed as directed in the kit. To measure cAMP concentration in zebrafish, fifteen 72 hpf zebrafish per condition were lysed in 175 *μ*L of 0.1M HCL lysis buffer. 100 *μ*M FSK treated zebrafish were used as a positive control.

### Statistics

All cardiac parameters from PV looping analysis are presented as means ± S.D. and SigmaStat 3.5 was used for comparison. A paired t test was used to compare data in same group before and after chemical infusion, while other data between groups were compared with one-way ANOVA. The significant difference between groups in dnPKA-related experiments were determined using two-way ANOVA. A post hoc Student-Newman-Keuls test was further conducted to compare difference between two groups. A P value<0.05 was considered significantly.

## Data availability

All data generated and analyzed during this study are included in this manuscript as figures, and supplementary figures and table. Transgenic animals are readily available upon request.

## Acknowledgments

We thank members of the Irannejad lab and Dr. Larsen for assistance, advice and valuable discussion. We also thank members of the Reiter lab, particularly Drs. Melissa Truong and Semil Choksi for help with transgenic zebrafish experiments and the zebrafish facility at the cardiovascular research institute, UCSF. We also thanks Dr. Di Lang for help with calcium assay. These studies were supported by the National Institute on General Medicine (GM133521) to R.I and T32 training grant (HL007731-30) to T.L.

