## Supplementary Figures for "Cardiac contraction and relaxation are regulated by beta 1 adrenergic receptor-generated cAMP pools at distinct membrane locations"

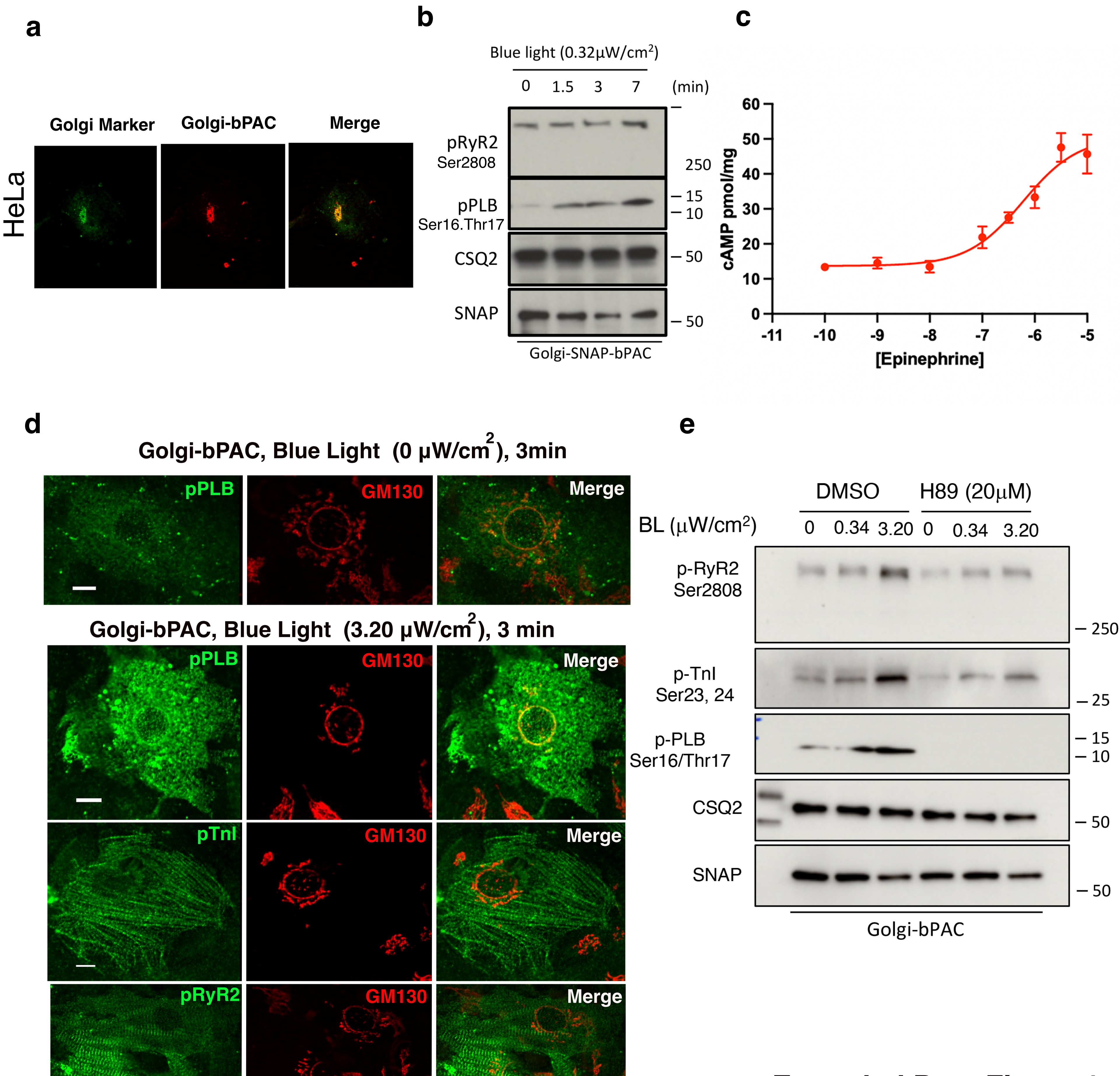


Extended Data Figure 1

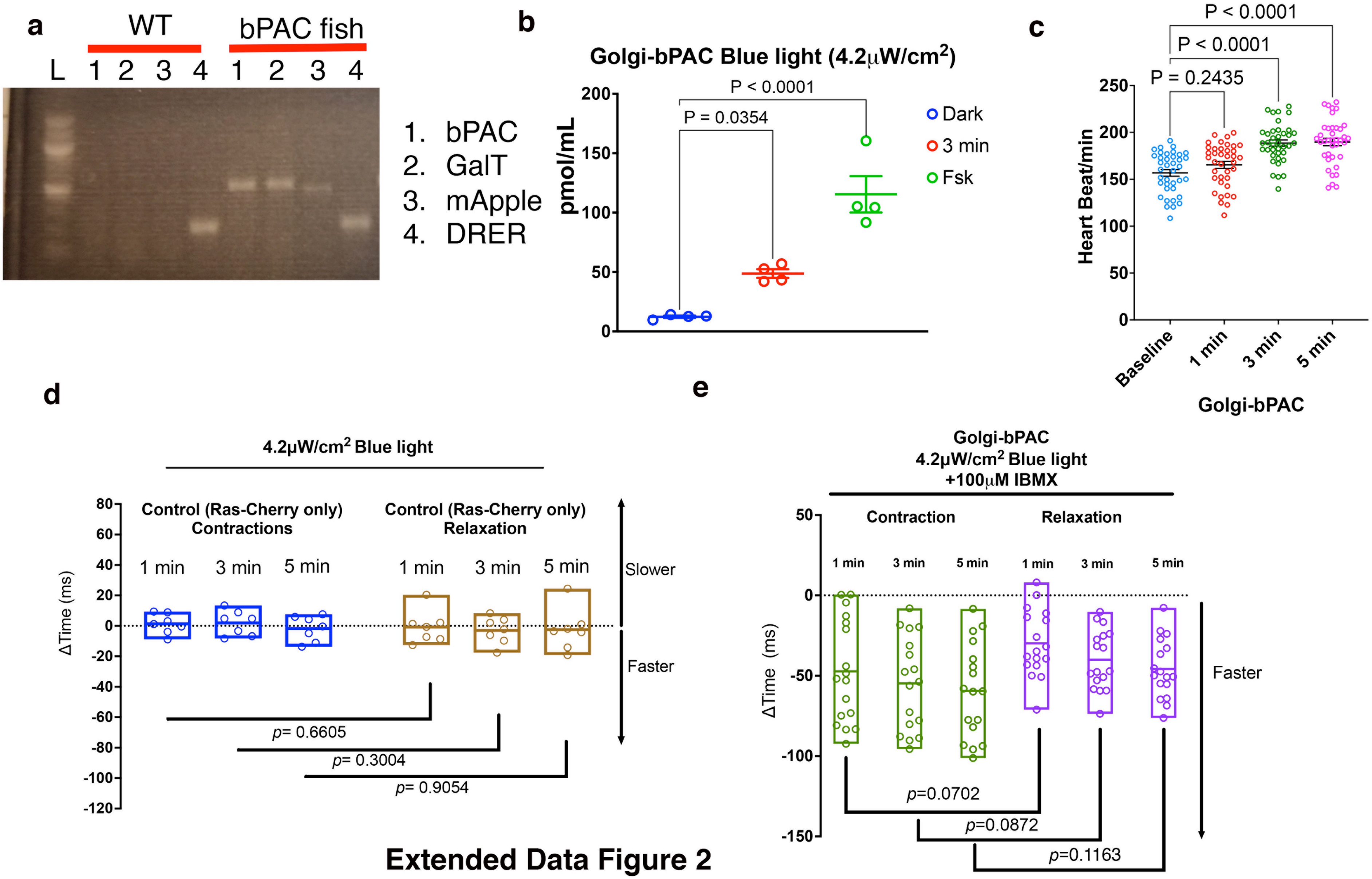


**Extended Data Figure 2**

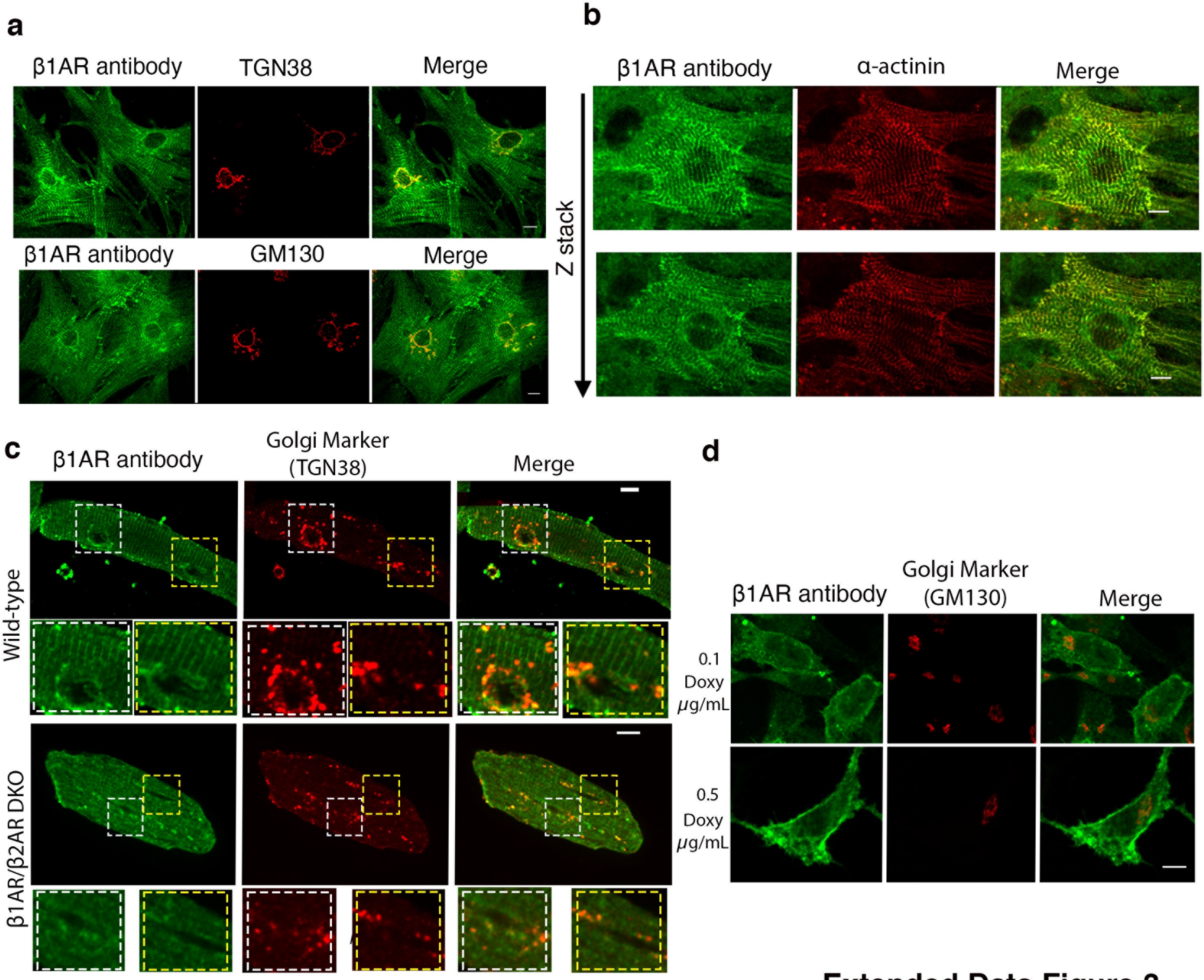


**Extended Data Figure 3**

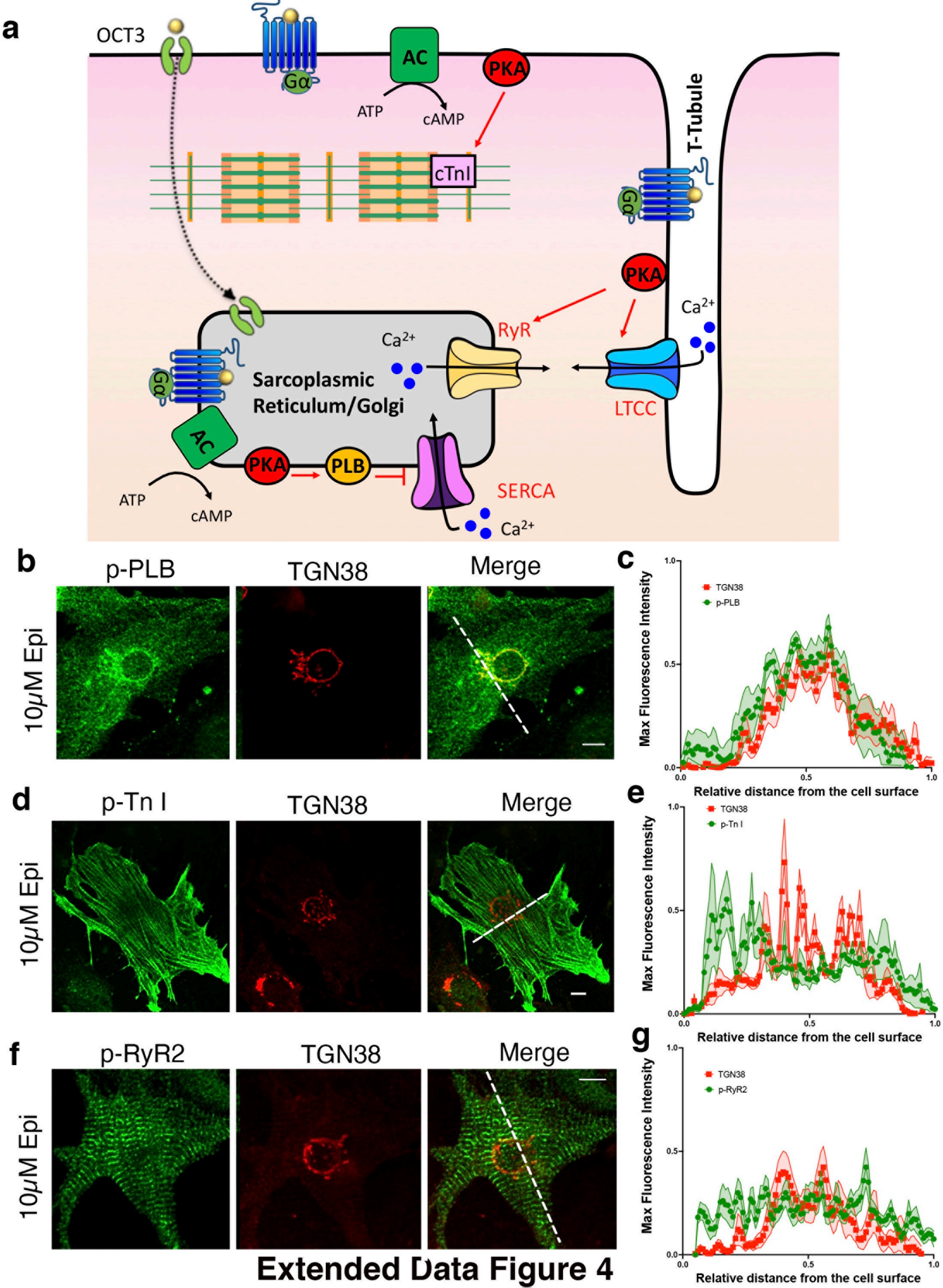


**Extended Data Figure 4**

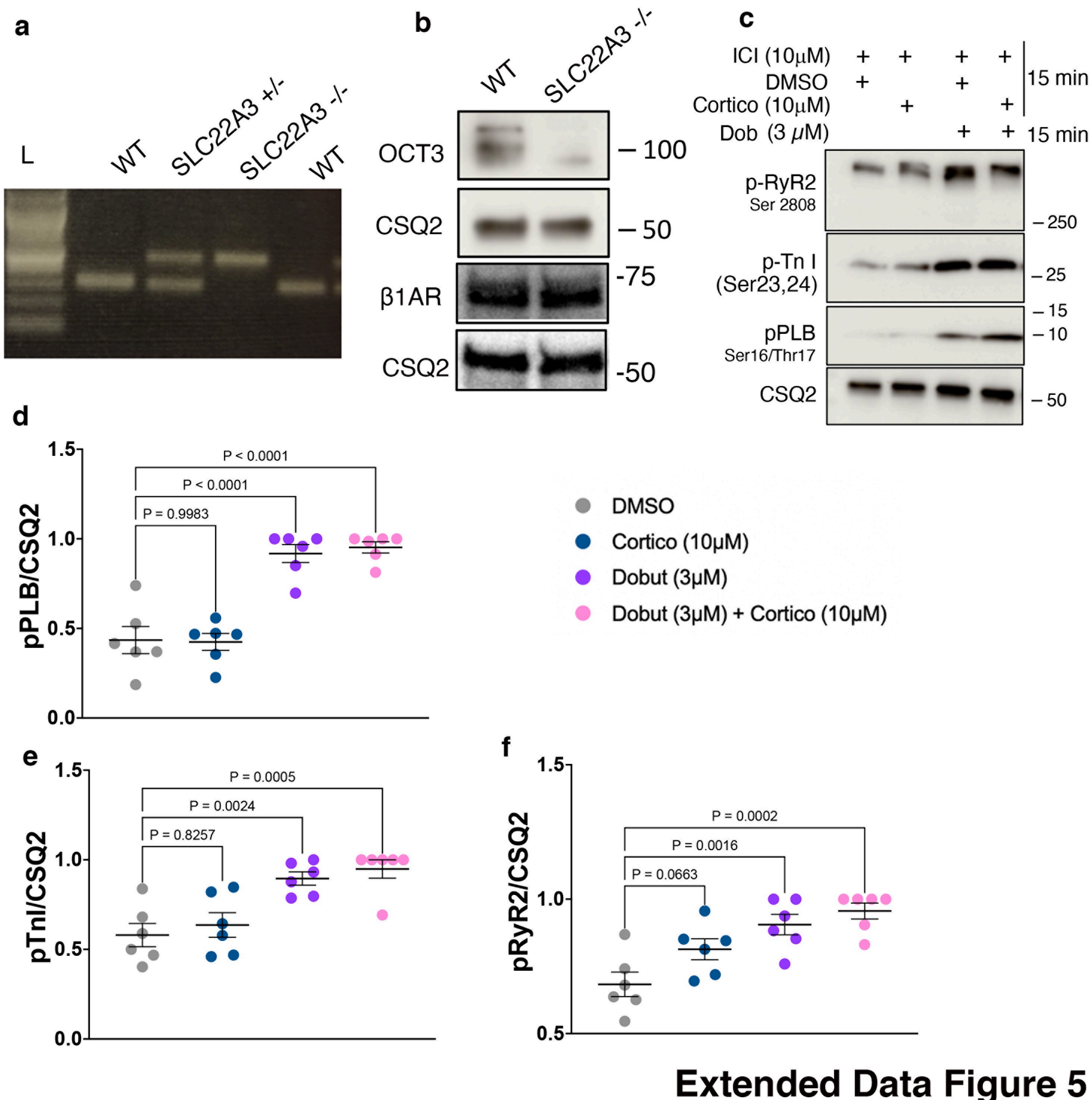


**Extended Data Figure 5**

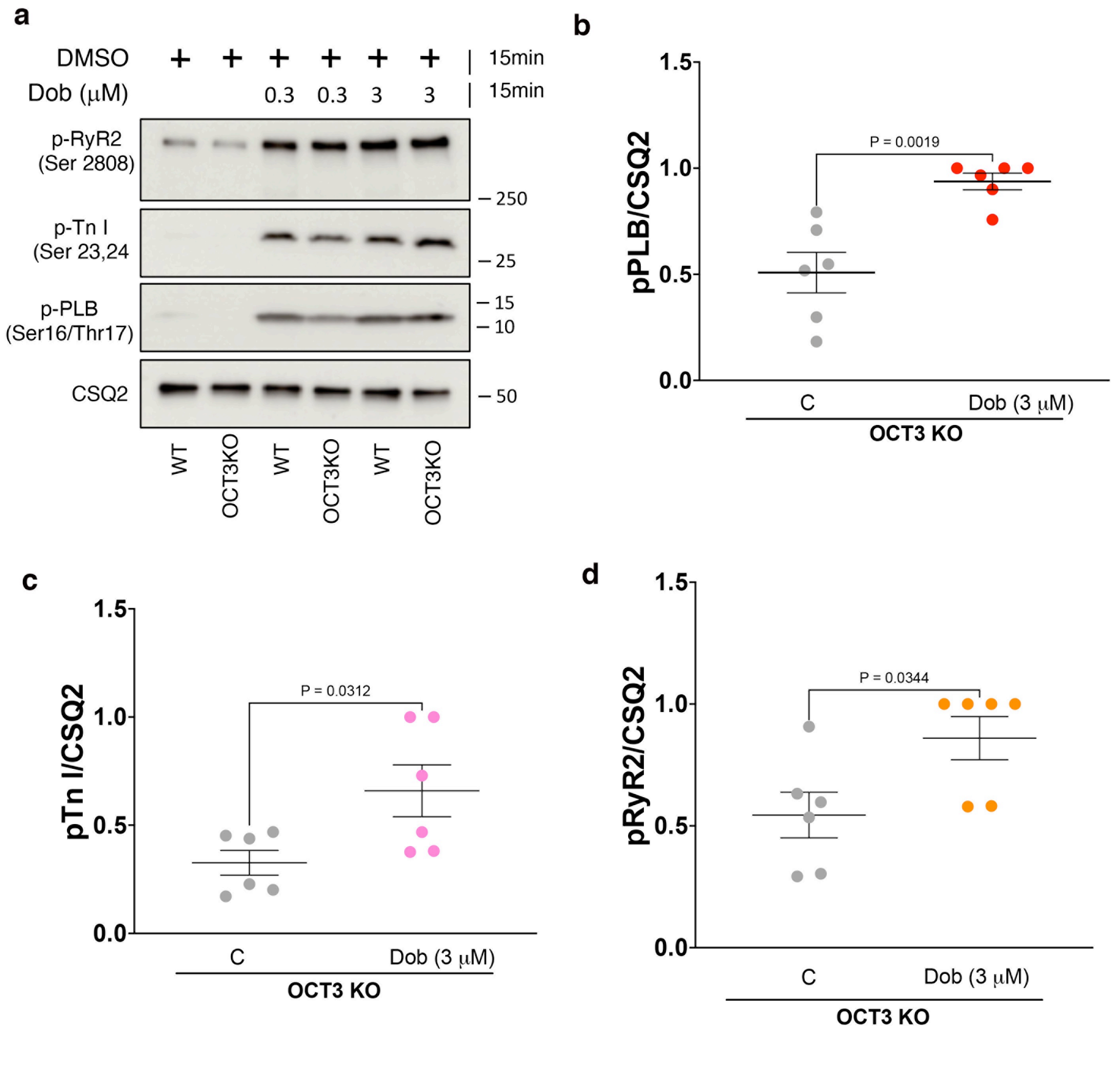


**Extended Data Figure 6**

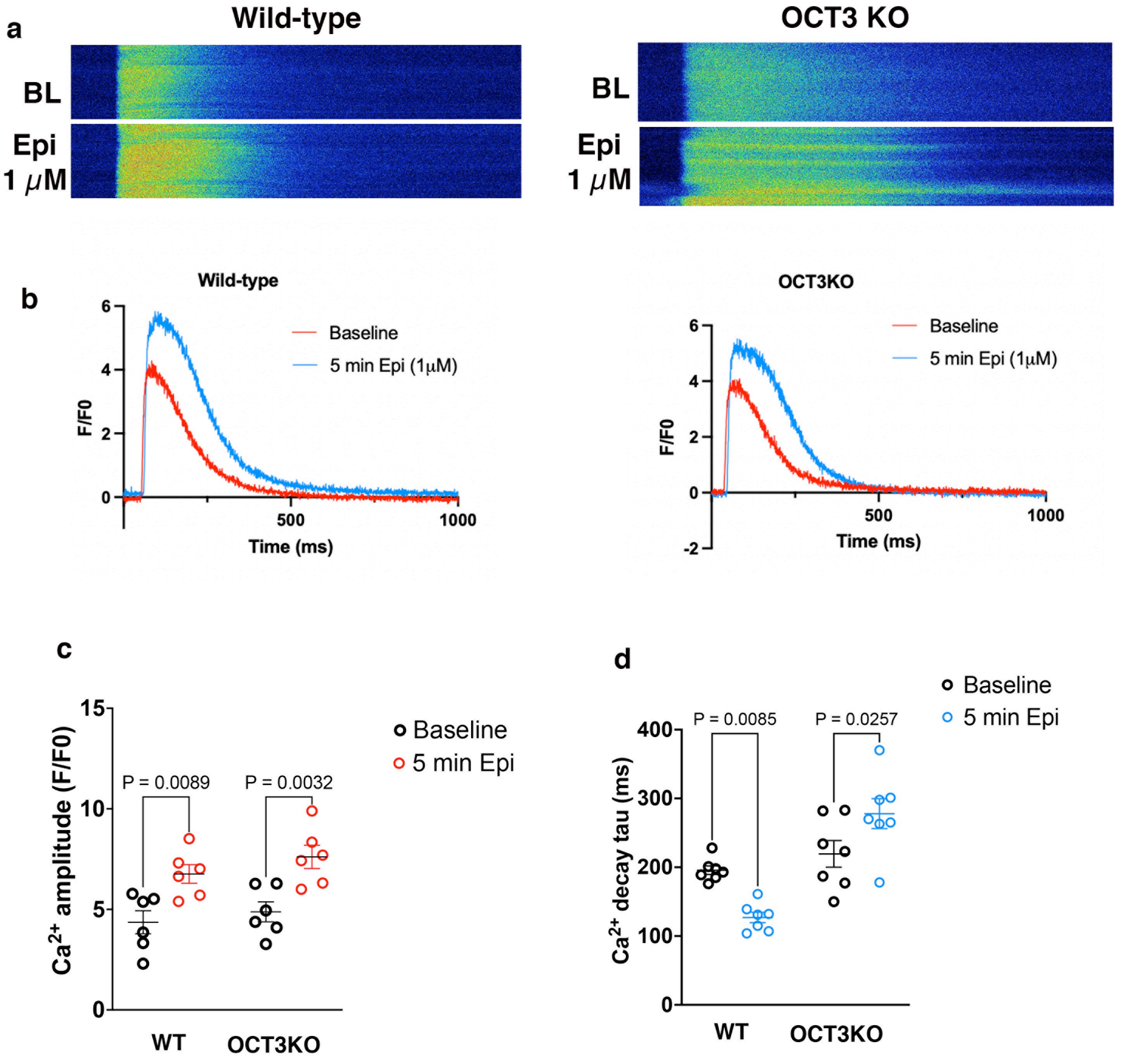


**Extended Data Figure 7**

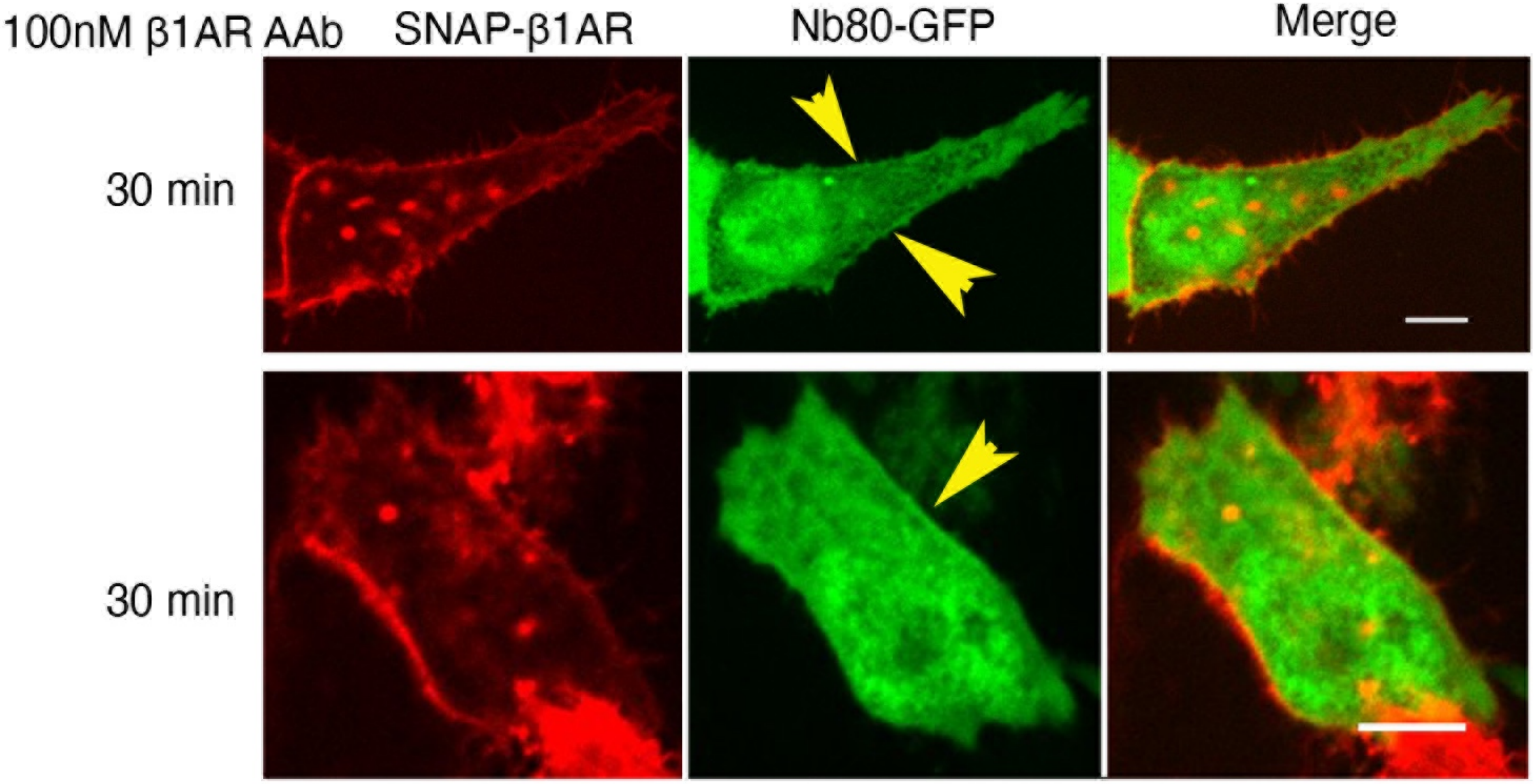


**Extended Data Figure 8**
